# The last bacterial common ancestor encoded the components of a functionally complete flagellum

**DOI:** 10.64898/2026.06.11.731707

**Authors:** Berkay Selçuk, Ekaterina P. Andrianova, Morgan Beeby, Daniel B. Kearns, Marc Erhardt, Igor B. Zhulin

## Abstract

Bacterial flagella are rotary nanomachines that enable motility in diverse environments. Although more than 40 genes are required for a functional flagellum in model organisms, only 24 have previously been inferred to be ancestral. Did the last bacterial common ancestor (LBCA) encode a simpler flagellum that was subsequently elaborated, or has its complexity been underestimated because highly diverged homologs are difficult to detect? Here, we combine homology searches across 80,789 bacterial proteomes with conserved sequence signatures, gene-tree clustering, and gene-neighborhood evidence to reconstruct the ancestral flagellum. We identify 49 flagellar protein groups with high-confidence support for ancestral presence across three phylogenetic reconstructions and one additional likely ancestral component, yielding a reconstructed repertoire of 50 ancestral protein groups. This repertoire includes the major structural, export, motor, and assembly components required for a functional flagellum. We provide an interactive database (https://www.flagelladb.org) for exploring the distribution, co-presence, and phylogenetic relationships of flagellar components across bacteria. Our findings indicate that the LBCA possessed a complex, functionally complete flagellar system, substantially exceeding previous estimates of ancestral complexity. The distribution of these components among extant bacteria is consistent with extensive lineage-specific loss and remodeling during subsequent flagellar evolution.

## INTRODUCTION

The bacterial flagellum is one of the best-studied molecular machines ^1,2^. It drives motility and chemotaxis ^3^, influences colony formation ^4^ and biofilm development ^5^, and plays key roles in pathogenicity ^6^ and host colonization ^7,8^. Flagellar motility is widespread among bacteria ^9^, and despite its high energetic cost, flagella are retained across diverse lineages, indicating a substantial contribution to fitness. In model organisms, it has been characterized in exceptional detail at the genetic, structural, and functional levels. Its biogenesis and assembly are tightly regulated, with components added in a precise temporal and spatial order, and it is integrated with complex signal transduction pathways ^10–12^. Owing to this combination of structural complexity and multi-step regulation, the flagellum is often cited as a model for understanding complex molecular systems ^13^. As such, it provides a powerful framework for investigating how complex molecular machines evolve, which remains a central question in biology.

The flagellum is composed of dozens of proteins organized into substructures, including basal body, hook, filament, and export apparatus, that together form a functional rotary nanomachine ^10–12^. Assembly begins with the export apparatus in the inner membrane, around which the MS-ring forms and provides the foundation for the cytoplasmic rotor (C-ring) and periplasmic driveshaft (rod) ^14–16^. The rod and subsequent axial structures are built from subunits transported by the export apparatus, powered by the proton motive force and ATP hydrolysis ^17^. Additional proteins form channels through the peptidoglycan and outer membrane, allowing the rod to extend beyond the cell envelope ^18–20^. A flexible universal joint (the hook) then forms at the rod tip, and upon reaching its characteristic length, a molecular ruler triggers an export-specificity switch that initiates construction of the filament, which can extend up to 10 μm ^21–23^. Finally, stator complexes assemble around the rotor, where ion flux generates torque that drives flagellar rotation ^24,25^. In addition to these structural components, regulatory proteins, chaperones, and assembly factors coordinate flagellar gene expression and the ordered assembly of the flagellum ^11,12^.

The prevailing view is that many bacteria share a conserved core set of flagellar genes supplemented by lineage-specific components ^26^. Previous comparative studies identified 24 core flagellar genes that were likely present in the last bacterial common ancestor (LBCA) ^26,27^. More recent reconstruction of the bacterial ancestor on a rooted bacterial phylogeny provided strong support for a flagellated LBCA ^28^, although the composition of the ancestral flagellar system was not systematically resolved. Phylogenetic analyses have documented repeated transitions in flagellar motility during bacterial diversification ^29,30^. In particular, Philip et al. ^30^ found that losses substantially exceeded gains for many flagellar proteins, suggesting an important role for secondary loss in shaping contemporary flagellar repertoires. Yet experimentally studied flagellar systems typically require more than 40 genes for their assembly, function and regulation ^11,31,32^. Notably, the highly reduced genome of *Buchnera aphidicola* retains a reduced set of flagellar genes and can assemble flagellar basal bodies but not a complete, functional flagellum ^33^. Thus, a substantial gap remains between the relatively small ancestral core inferred from earlier comparative analyses and the complexity of modern flagellar systems, in part because highly diverged homologs can be difficult to detect across deep evolutionary timescales ^27,34,35^. This raises a fundamental question about flagellar evolution: did the LBCA possess a simpler flagellum that was subsequently elaborated, or has the ancestral system been underestimated?

Here, using a comprehensive computational framework, we identify orthologous groups of flagellar proteins across bacteria, defined as sets of proteins likely to share the same function that may include both orthologs and lineage-specific paralogs. By combining qualitative assessment of phyletic gene distribution and ancestral state reconstruction, we infer the flagellar gene repertoire of the LBCA and show that it was substantially more complete than previously recognized, closely resembling that of extant bacteria and indicating that a functionally complete, near-modern flagellum was already present in the LBCA.

## RESULTS

### Defining a non-redundant flagellar protein dataset

To compile a comprehensive set of flagellum-associated proteins, we used the KEGG pathway map02040 (“Flagellar assembly”), which lists 54 proteins, as an initial reference set. To identify additional conserved flagellum-associated genes that may not be represented in KEGG, we compiled a representative set of 251 genomes of flagellated bacterial species. Genomes were selected to provide broad taxonomic coverage, include experimentally studied organisms, and prioritize genome quality and representation in the MiST database ^36^. Comparison of representative experimentally characterized systems illustrates both the conservation of major flagellar modules and substantial variation in their overall protein complements (Fig. 1).

**Figure 1:**
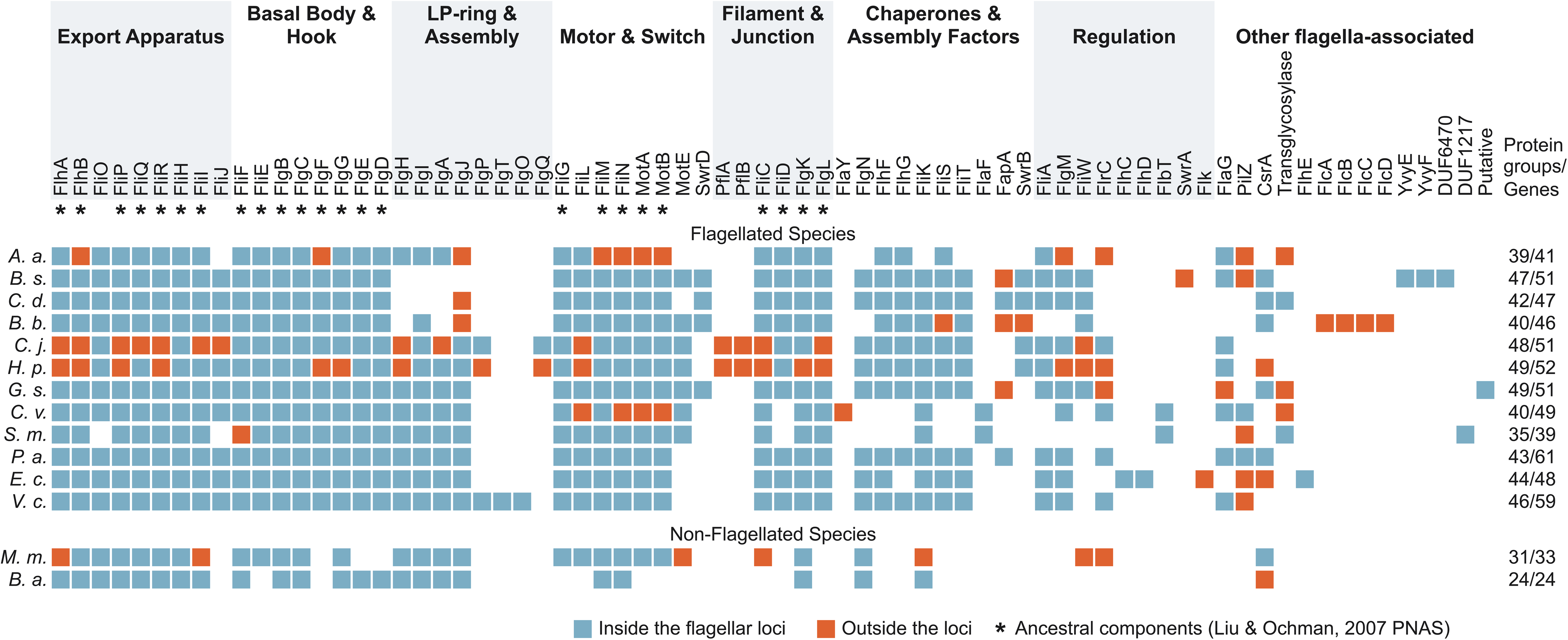
Diversity of flagellar protein complements in representative bacterial species. Distribution of flagellum-associated protein groups (squares) across representative bacterial species selected to illustrate diversity of flagellar gene complements. Proteins are organized according to their principal roles in flagellar assembly, structure, function, and regulation. Protein groups previously proposed to constitute the ancestral flagellar core ^26^ are marked by asterisks. Blue squares indicate proteins encoded within flagellar loci together with other flagellar components, whereas orange squares indicate flagellum-associated proteins encoded outside these loci. The proteins shown include both previously characterized flagellar components and additional proteins identified in this study through their association with flagellar loci. Numbers on the right indicate the number of distinct flagellum-associated protein groups followed by the total number of flagellar genes. For example, the *A. aeolicus* genome contains 39 distinct flagellum-associated protein groups and additional copies of *motB* and *pilZ*, for a total of 41 flagellar genes. Species abbreviations (genome accessions): A.a., *Aquifex aeolicus* (GCF_000008625.1); B.s., *Bacillus subtilis* (GCF_002153395.1); C.d., *Clostridioides difficile* (GCF_000438845.1); B.b., *Borreliella burgdorferi* (GCF_000008685.2); C.j., *Campylobacter jejuni* (GCF_001457695.1); H.p., *Helicobacter pylori* (GCF_900478295.1); G.s., *Geobacter sulfurreducens* (GCF_000007985.2); C.v., *Caulobacter vibrioides* (GCF_002858865.1); S.m., *Sinorhizobium meliloti* (GCF_000375585.1); P.a., *Pseudomonas aeruginosa* (GCF_001457615.1); E.c., *Escherichia coli* (GCF_003697165.2); V.c., *Vibrio cholerae* (GCF_001471585.2); M.m., *Midichloria mitochondrii* (GCF_000219355.1); B.a., *Buchnera aphidicola* (GCF_000225445.1).

We searched the complete set of 251 genomes using profile hidden Markov models (HMMs) for each of the 54 flagellar proteins (see Methods). The distribution of these proteins across the representative genomes is provided in Table S1. Inspection of flagellar gene neighborhoods identified several additional genes that were consistently associated with flagellar loci across diverse bacteria including *yvyF* and *yviE.* Each was identified in 32 genomes (∼13% of the representative dataset), and in every instance the gene was located within or immediately adjacent to a flagellar gene cluster. This consistent association justified their inclusion in the set of flagellum-associated proteins. Chemotaxis genes were also frequently found within or adjacent to flagellar loci. However, because they encode components of a distinct, well-characterized signal-transduction system rather than flagella, they were not included in our analysis. We further expanded this candidate set through an extensive literature survey focused primarily on experimentally characterized genes in model organisms. Together, these approaches yielded an initial list of 95 proteins involved in the assembly, function, and regulation of flagella in diverse bacterial species. Following manual curation, synonymous gene names and homologous proteins representing the same functional group were consolidated into unified orthologous groups (Table 1). The resulting non-redundant dataset comprised 81 orthologous groups of known and putative flagellum-associated proteins (Table 2) which served as the reference set for downstream large-scale homology searches.

**Table 1.**
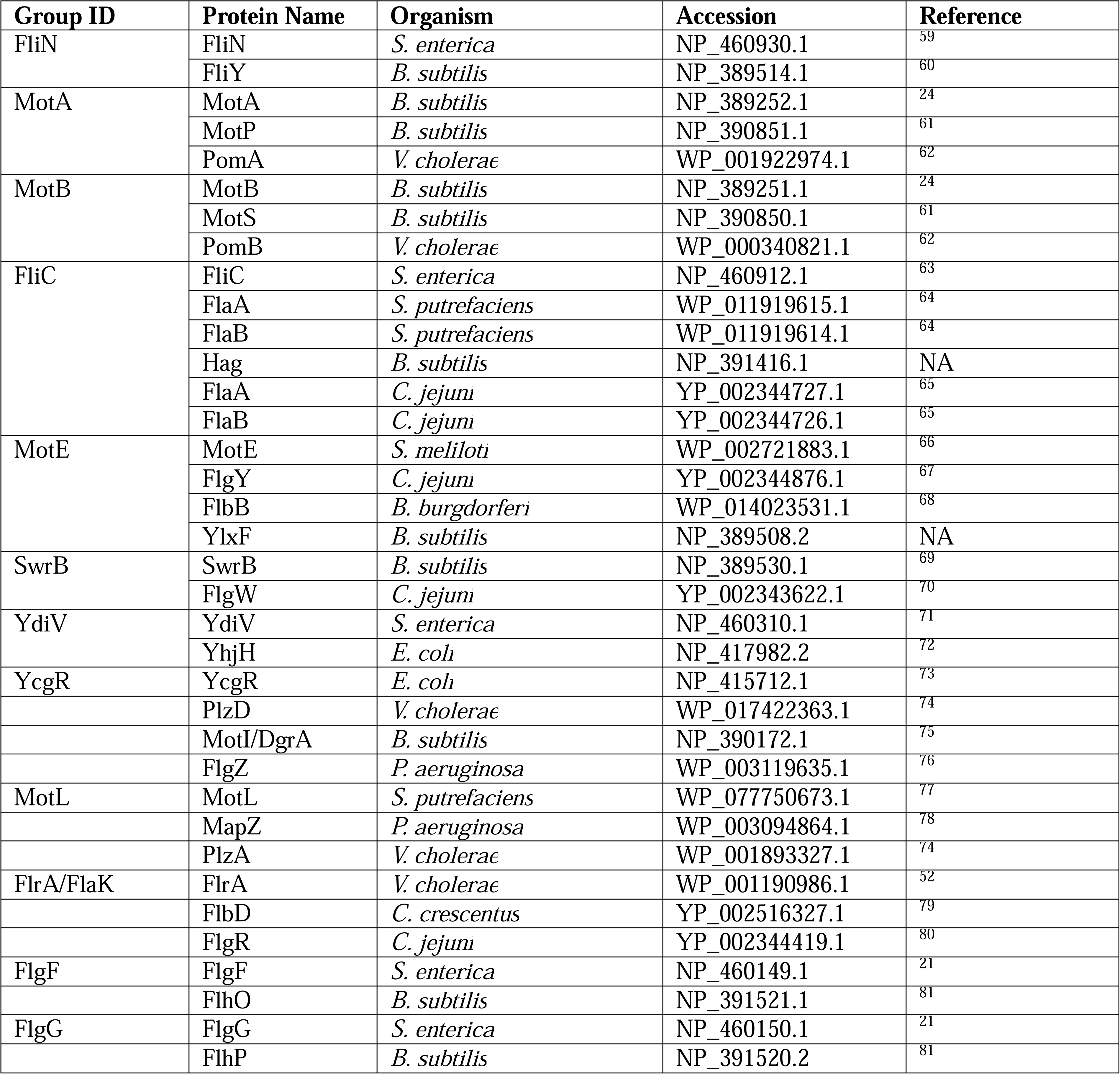
Orthologous groups of flagellar components with different names.

**Table 2.**
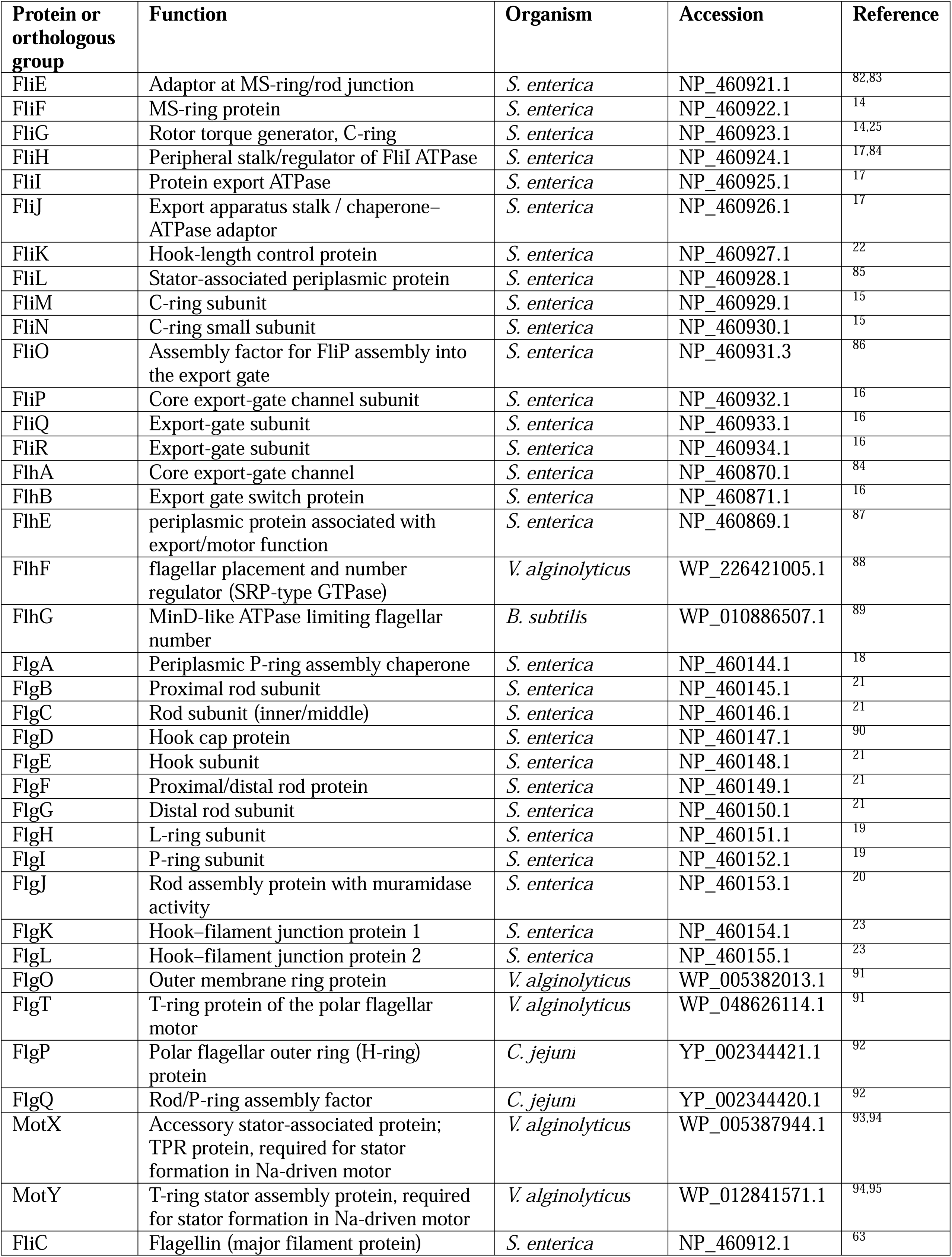
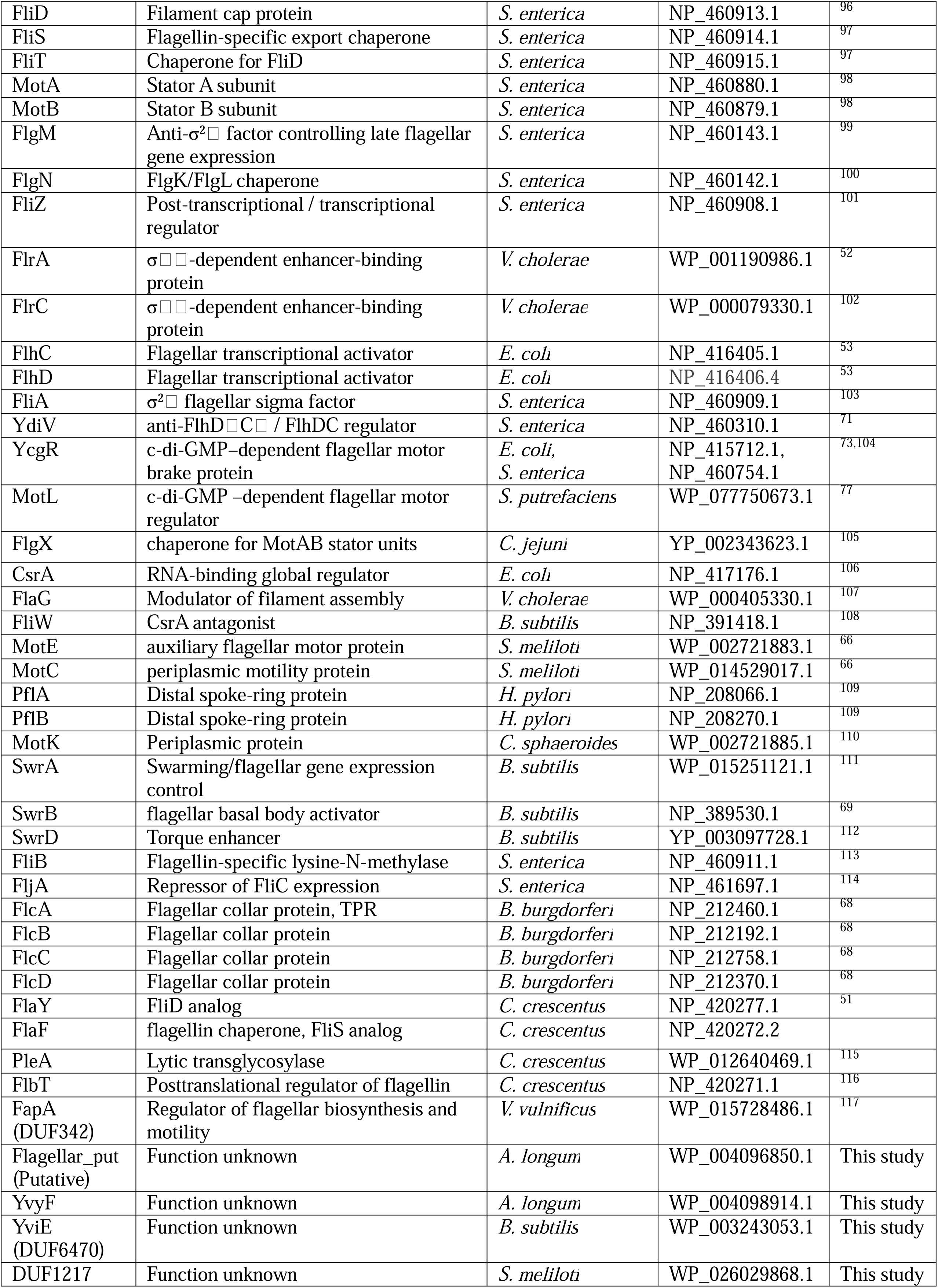
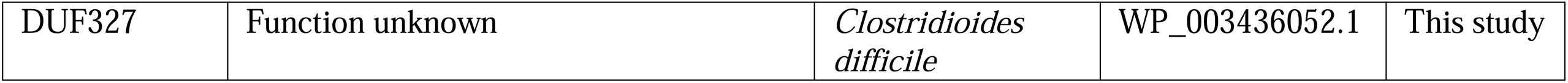
Proteins involved in assembly, function, and regulation of bacterial flagella.

| Protein or orthologous group | Function | Organism | Accession | Reference |
|---|---|---|---|---|
| FliE | Adaptor at MS-ring/rod junction | *S. enterica* | NP_460921.1 | <sup>82,83</sup> |
| FliF | MS-ring protein | *S. enterica* | NP_460922.1 | <sup>14</sup> |
| FliG | Rotor torque generator, C-ring | *S. enterica* | NP_460923.1 | <sup>14,25</sup> |
| FliH | Peripheral stalk/regulator of FliI ATPase | *S. enterica* | NP_460924.1 | <sup>17,84</sup> |
| FliI | Protein export ATPase | *S. enterica* | NP_460925.1 | <sup>17</sup> |
| FliJ | Export apparatus stalk / chaperone–ATPase adaptor | *S. enterica* | NP_460926.1 | <sup>17</sup> |
| FliK | Hook-length control protein | *S. enterica* | NP_460927.1 | <sup>22</sup> |
| FliL | Stator-associated periplasmic protein | *S. enterica* | NP_460928.1 | <sup>85</sup> |
| FliM | C-ring subunit | *S. enterica* | NP_460929.1 | <sup>15</sup> |
| FliN | C-ring small subunit | *S. enterica* | NP_460930.1 | <sup>15</sup> |
| FliO | Assembly factor for FliP assembly into the export gate | *S. enterica* | NP_460931.3 | <sup>86</sup> |
| FliP | Core export-gate channel subunit | *S. enterica* | NP_460932.1 | <sup>16</sup> |
| FliQ | Export-gate subunit | *S. enterica* | NP_460933.1 | <sup>16</sup> |
| FliR | Export-gate subunit | *S. enterica* | NP_460934.1 | <sup>16</sup> |
| FlhA | Core export-gate channel | *S. enterica* | NP_460870.1 | <sup>84</sup> |
| FlhB | Export gate switch protein | *S. enterica* | NP_460871.1 | <sup>16</sup> |
| FlhE | periplasmic protein associated with export/motor function | *S. enterica* | NP_460869.1 | <sup>87</sup> |
| FlhF | flagellar placement and number regulator (SRP-type GTPase) | *V. alginolyticus* | WP_226421005.1 | <sup>88</sup> |
| FlhG | MinD-like ATPase limiting flagellar number | *B. subtilis* | WP_010886507.1 | <sup>89</sup> |
| FlgA | Periplasmic P-ring assembly chaperone | *S. enterica* | NP_460144.1 | <sup>18</sup> |
| FlgB | Proximal rod subunit | *S. enterica* | NP_460145.1 | <sup>21</sup> |
| FlgC | Rod subunit (inner/middle) | *S. enterica* | NP_460146.1 | <sup>21</sup> |
| FlgD | Hook cap protein | *S. enterica* | NP_460147.1 | <sup>90</sup> |
| FlgE | Hook subunit | *S. enterica* | NP_460148.1 | <sup>21</sup> |
| FlgF | Proximal/distal rod protein | *S. enterica* | NP_460149.1 | <sup>21</sup> |
| FlgG | Distal rod subunit | *S. enterica* | NP_460150.1 | <sup>21</sup> |
| FlgH | L-ring subunit | *S. enterica* | NP_460151.1 | <sup>19</sup> |
| FlgI | P-ring subunit | *S. enterica* | NP_460152.1 | <sup>19</sup> |
| FlgJ | Rod assembly protein with muramidase activity | *S. enterica* | NP_460153.1 | <sup>20</sup> |
| FlgK | Hook–filament junction protein 1 | *S. enterica* | NP_460154.1 | <sup>23</sup> |
| FlgL | Hook–filament junction protein 2 | *S. enterica* | NP_460155.1 | <sup>23</sup> |
| FlgO | Outer membrane ring protein | *V. alginolyticus* | WP_005382013.1 | <sup>91</sup> |
| FlgT | T-ring protein of the polar flagellar motor | *V. alginolyticus* | WP_048626114.1 | <sup>91</sup> |
| FlgP | Polar flagellar outer ring (H-ring) protein | *C. jejuni* | YP_002344421.1 | <sup>92</sup> |
| FlgQ | Rod/P-ring assembly factor | *C. jejuni* | YP_002344420.1 | <sup>92</sup> |
| MotX | Accessory stator-associated protein; TPR protein, required for stator formation in Na-driven motor | *V. alginolyticus* | WP_005387944.1 | <sup>93,94</sup> |
| MotY | T-ring stator assembly protein, required for stator formation in Na-driven motor | *V. alginolyticus* | WP_012841571.1 | <sup>94,95</sup> |
| FliC | Flagellin (major filament protein) | *S. enterica* | NP_460912.1 | <sup>63</sup> |
| FliD | Filament cap protein | *S. enterica* | NP_460913.1 | <sup>96</sup> |
| FliS | Flagellin-specific export chaperone | *S. enterica* | NP_460914.1 | <sup>97</sup> |
| FliT | Chaperone for FliD | *S. enterica* | NP_460915.1 | <sup>97</sup> |
| MotA | Stator A subunit | *S. enterica* | NP_460880.1 | <sup>98</sup> |
| MotB | Stator B subunit | *S. enterica* | NP_460879.1 | <sup>98</sup> |
| FlgM | Anti-σ²□ factor controlling late flagellar gene expression | *S. enterica* | NP_460143.1 | <sup>99</sup> |
| FlgN | FlgK/FlgL chaperone | *S. enterica* | NP_460142.1 | <sup>100</sup> |
| FliZ | Post-transcriptional / transcriptional regulator | *S. enterica* | NP_460908.1 | <sup>101</sup> |
| FlrA | σ□□-dependent enhancer-binding protein | *V. cholerae* | WP_001190986.1 | <sup>52</sup> |
| FlrC | σ□□-dependent enhancer-binding protein | *V. cholerae* | WP_000079330.1 | <sup>102</sup> |
| FlhC | Flagellar transcriptional activator | *E. coli* | NP_416405.1 | <sup>53</sup> |
| FlhD | Flagellar transcriptional activator | *E. coli* | NP_416406.4 | <sup>53</sup> |
| FliA | σ²□ flagellar sigma factor | *S. enterica* | NP_460909.1 | <sup>103</sup> |
| YdiV | anti-FlhD□C□ / FlhDC regulator | *S. enterica* | NP_460310.1 | <sup>71</sup> |
| YcgR | c-di-GMP–dependent flagellar motor brake protein | *E. coli, S. enterica* | NP_415712.1, NP_460754.1 | <sup>73,104</sup> |
| MotL | c-di-GMP –dependent flagellar motor regulator | *S. putrefaciens* | WP_077750673.1 | <sup>77</sup> |
| FlgX | chaperone for MotAB stator units | *C. jejuni* | YP_002343623.1 | <sup>105</sup> |
| CsrA | RNA-binding global regulator | *E. coli* | NP_417176.1 | <sup>106</sup> |
| FlaG | Modulator of filament assembly | *V. cholerae* | WP_000405330.1 | <sup>107</sup> |
| FliW | CsrA antagonist | *B. subtilis* | NP_391418.1 | <sup>108</sup> |
| MotE | auxiliary flagellar motor protein | *S. meliloti* | WP_002721883.1 | <sup>66</sup> |
| MotC | periplasmic motility protein | *S. meliloti* | WP_014529017.1 | <sup>66</sup> |
| PflA | Distal spoke-ring protein | *H. pylori* | NP_208066.1 | <sup>109</sup> |
| PflB | Distal spoke-ring protein | *H. pylori* | NP_208270.1 | <sup>109</sup> |
| MotK | Periplasmic protein | *C. sphaeroides* | WP_002721885.1 | <sup>110</sup> |
| SwrA | Swarming/flagellar gene expression control | *B. subtilis* | WP_015251121.1 | <sup>111</sup> |
| SwrB | flagellar basal body activator | *B. subtilis* | NP_389530.1 | <sup>69</sup> |
| SwrD | Torque enhancer | *B. subtilis* | YP_003097728.1 | <sup>112</sup> |
| FliB | Flagellin-specific lysine-N-methylase | *S. enterica* | NP_460911.1 | <sup>113</sup> |
| FljA | Repressor of FliC expression | *S. enterica* | NP_461697.1 | <sup>114</sup> |
| FlcA | Flagellar collar protein, TPR | *B. burgdorferi* | NP_212460.1 | <sup>68</sup> |
| FlcB | Flagellar collar protein | *B. burgdorferi* | NP_212192.1 | <sup>68</sup> |
| FlcC | Flagellar collar protein | *B. burgdorferi* | NP_212758.1 | <sup>68</sup> |
| FlcD | Flagellar collar protein | *B. burgdorferi* | NP_212370.1 | <sup>68</sup> |
| FlaY | FliD analog | *C. crescentus* | NP_420277.1 | <sup>51</sup> |
| FlaF | flagellin chaperone, FliS analog | *C. crescentus* | NP_420272.2 |  |
| PleA | Lytic transglycosylase | *C. crescentus* | WP_012640469.1 | <sup>115</sup> |
| FlbT | Posttranslational regulator of flagellin | *C. crescentus* | NP_420271.1 | <sup>116</sup> |
| FapA (DUF342) | Regulator of flagellar biosynthesis and motility | *V. vulnificus* | WP_015728486.1 | <sup>117</sup> |
| Flagellar_put (Putative) | Function unknown | *A. longum* | WP_004096850.1 | This study |
| YvyF | Function unknown | *A. longum* | WP_004098914.1 | This study |
| YviE (DUF6470) | Function unknown | *B. subtilis* | WP_003243053.1 | This study |
| DUF1217 | Function unknown | *S. meliloti* | WP_026029868.1 | This study |
| DUF327 | Function unknown | *Clostridioides difficile* | WP_003436052.1 | This study |

### Identifying flagellar proteins across the bacterial tree of life

To identify flagellar proteins across bacteria, we developed a computational framework (Fig. 2) for sensitive detection of orthologous groups across GTDB release 214 ^37^, comprising 80,789 representative bacterial proteomes. For each of the 81 curated flagellum-associated protein groups, we used Hidden Markov Models (HMMs) from the KOfamKOALA ^38^, Pfam ^39^, TIGRFAMs ^40^, and NCBIfam ^41^ databases where available, or representative annotated protein sequences as queries for iterative profile searches otherwise. Searches were performed using relaxed E-value thresholds to maximize sensitivity and capture divergent homologs. For most proteins, we retained the top 50,000 hits, a cutoff chosen empirically to capture a large collection of candidate sequences (given that about one third of the ∼80,000 genomes were expected to encode flagella ^35^), while keeping subsequent analyses computationally tractable. For multicopy proteins such as FliC, MotA, and MotB, we retained the top 100,000 hits because their large orthologous groups required deeper retrieval both to capture most members and to include sufficient background sequences for delineating group boundaries.

**Figure 2:**
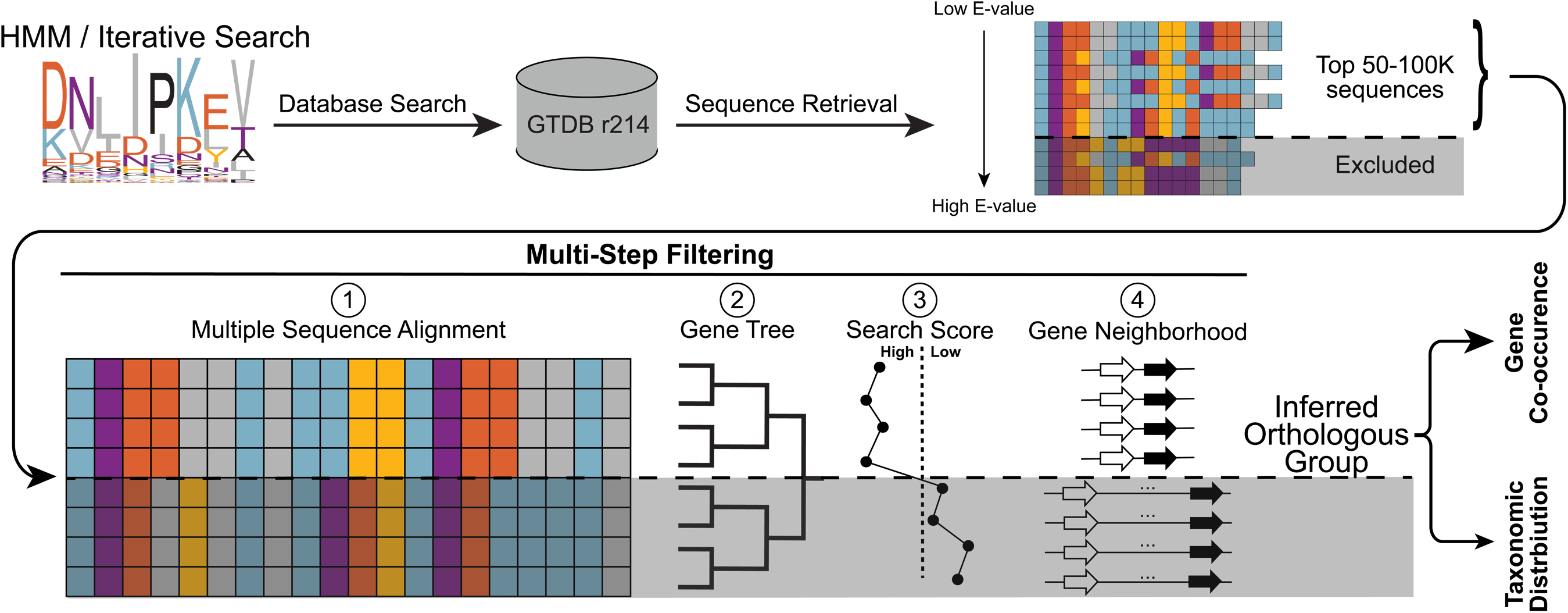
Computational framework for identifying flagellar proteins across Bacteria. For each flagellar protein, the corresponding HMMs or representative protein sequences were used to search for homologous sequences against the GTDB r214 database. The top 50-100K hits, ranked by their search score, were retrieved, aligned, and used to construct gene trees. Orthologous groups of each flagellar protein were then inferred using four complementary criteria: conserved sequence patterns in multiple sequence alignments, clustering within gene trees, higher search scores, and co-localization with other flagellar genes within the same genomic neighborhood.

Sequences within clades satisfying the combined identification criteria were retained as members of the corresponding orthologous groups for downstream analyses. These criteria included phylogenetic clustering, sequence similarity scores, conserved alignment patterns, and genomic neighborhood associations with other flagellar genes (Fig. 2, Supplementary Fig. 1). Importantly, the criteria were evaluated collectively at the clade level rather than applied as strict filters to individual sequences. Consequently, sequences with relatively low similarity scores, or without association with flagellar gene neighborhoods could still be retained when supported by the combined evidence. Here, we use the term *orthologous group* consistent with the conventions established by databases such as COG ^42^ and eggNOG ^43^ to describe evolutionarily related proteins corresponding to individual flagellar components. Thus, each orthologous group may include vertically inherited orthologs, paralogs arising from gene duplication, and xenologs acquired via horizontal gene transfer.

To estimate the sensitivity of our bacteria-wide detection pipeline, we compared its results with the manually curated reference dataset using the 238 genome assemblies shared between the two datasets and 80 matched protein groups (Table S2). Because neither dataset constitutes a definitive gold standard, we calculated true-positive rates in both directions. Using the manually curated set as the reference, the pipeline recovered a median of 98.7% of protein accessions per group. Conversely, using the pipeline-generated dataset as the reference, the manually curated dataset contained a median of 94.9% of pipeline-detected accessions. This difference may reflect additional divergent homologs recovered by the pipeline, false-positive assignments or both. Although this comparison cannot establish absolute sensitivity or specificity, the high concordance between the two independently assembled datasets supports the overall reliability of our protein assignments.

### Co-presence analysis reveals conserved and functionally coherent flagellar gene clusters

Following the identification of flagellar orthologous groups, we quantified pairwise gene co-presence across bacterial genomes to identify flagellar genes that are preferentially retained together. The rationale was that genes encoding components of the same functional module, and especially those required for a functional flagellum, might exhibit similar patterns of retention and loss across genomes. We therefore analyzed 55 widely distributed flagellar protein groups and quantified pairwise co-presence using the Jaccard similarity index, defined as the fraction of genomes containing both genes among those containing at least one of the two (Fig. 3a). To account for taxonomic sampling bias, each genome was weighted inversely to the number of representative genomes in its taxonomic family, yielding a balanced weighted Jaccard similarity (J_w_), so that every family ends up contributing equally to the calculation regardless of how heavily it was sampled (Fig. 3b). Weighted similarities were converted to distances (1 − J_w_) and used for complete-linkage clustering (Fig. 3c).

**Figure 3:**
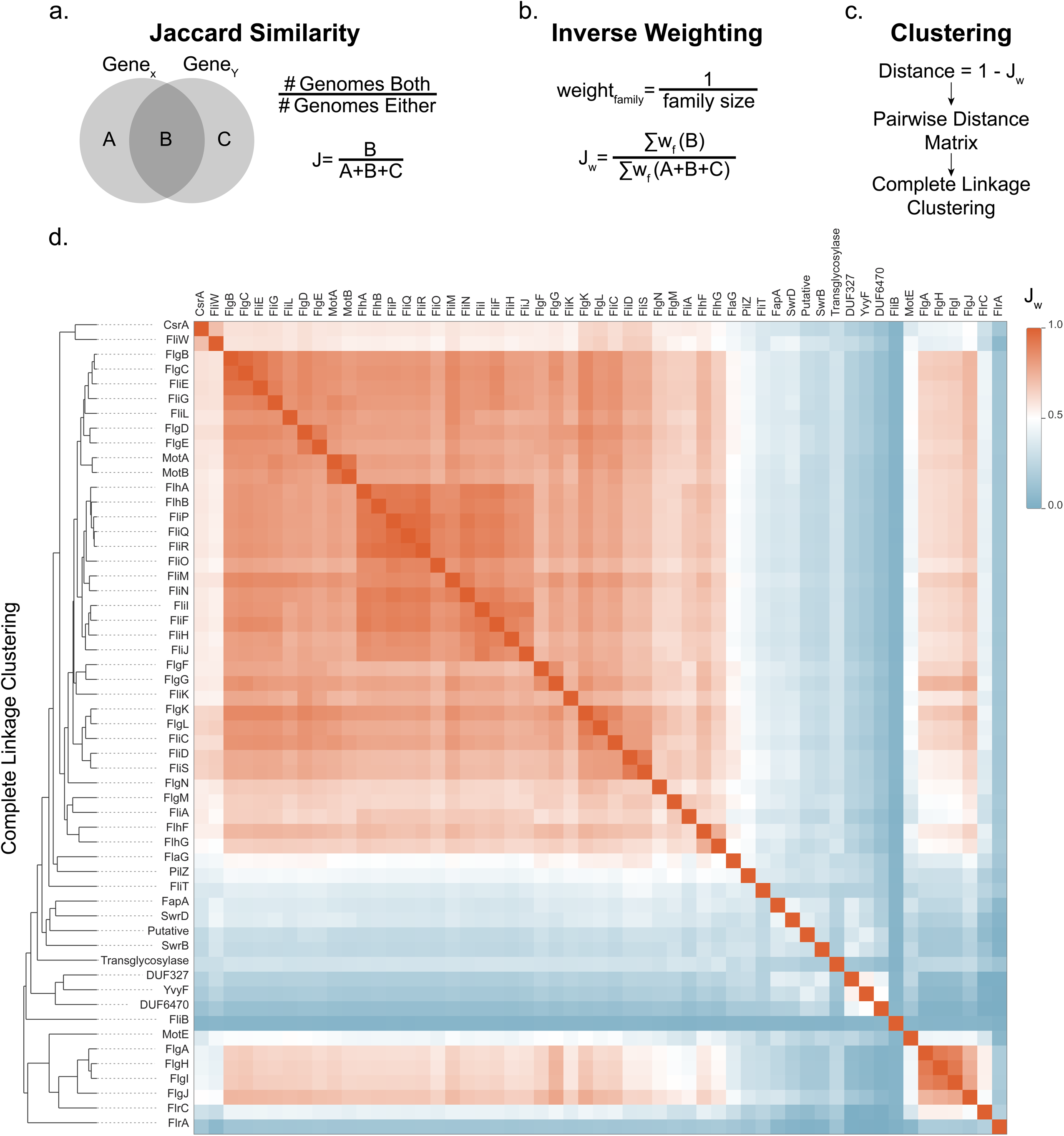
Pairwise co-presence analysis identifies conserved and auxiliary flagellar gene clusters. **a,** Schematic representation of the Jaccard similarity index, calculated as the fraction of genomes in which two genes co-present relative to the number of genomes containing at least one of the two genes. **b,** To correct for uneven taxonomic sampling, each genome was weighted inversely to the number of representative genomes in its family. The sums run over individual genomes, so that J_w_ is the summed weight of genomes encoding both genes (B) divided by the summed weight of genomes encoding at least one of them (A + B + C), and all families contribute equally. **c,** Weighted Jaccard similarities were converted to distances (1 − J_w_) and used for hierarchical clustering with complete linkage. **d,** Heatmap of pairwise family-weighted Jaccard similarities for 55 widely distributed flagellar protein groups across GTDB representative bacterial proteomes (release 214). Genes are ordered by complete-linkage hierarchical clustering, shown as a dendrogram on the left. The heatmap resolves broadly conserved core flagellar genes as well as more weakly associated auxiliary components. The tool that was used to create this plot is available at https://www.flagelladb.org/gene-correlation.

This analysis identified two major clusters of strongly co-present genes (J_w_ generally > 0.5). The larger cluster comprised 36 genes ranging from CsrA to FlhG in Fig. 3d, whereas the smaller cluster contained four genes from FlgA to FlgJ. Although separated by hierarchical clustering, these groups also showed strong cross-cluster similarity, indicating that they are frequently retained together across bacterial genomes. The smaller cluster likely separates because it contains proteins associated with the outer membrane and peptidoglycan layer (P and L rings) ^18–20^, which are frequently co-absent in monoderm genomes, likely reflecting differences in cell-envelope architecture ^44^. Together, these two clusters define a set of 40 strongly co-present flagellar genes, consistent with a conserved functional core and highlighting them as candidate components of the ancestral flagellar gene set.

Beyond this broadly conserved core, co-presence analysis also recovered local clusters corresponding to known structural and functional associations. The most tightly correlated sub-cluster comprised 12 proteins with homologs in both flagellar and non-flagellar type III secretion systems (T3SS), which we refer to as the T3SS-related module (FliJ, FliI, FliF, FliH, FliN, FliM, FliO, FlhA, FlhB, FliQ, FliP, and FliR; Supplementary Fig. 2). This module includes the export apparatus together with the MS-ring protein FliF and the C-ring proteins FliM and FliN. Its tight clustering likely reflects this shared occurrence, as non-flagellar T3SS homologs were not explicitly excluded by our search pipeline. Because non-flagellar T3SS were shown to have evolved from the flagellum ^45^, their components share the evolutionary history of the corresponding flagellar proteins, and their inclusion is therefore unlikely to distort the inferred ancestral presence of these protein groups. Other clusters corresponded to well-defined structural modules, including the stator unit (MotA, MotB), proximal rod components (FlgB, FlgC, FliE), distal rod components (FlgF, FlgG), P and L rings (FlgA, FlgH, FlgI, FlgJ), hook (FlgD, FlgE), hook–filament junction and filament (FlgK, FlgL, FliC), filament cap and chaperone of the filament (FliD, FliS). Regulatory proteins also clustered according to known functional relationships, including FliA/FlgM, FlhF/FlhG, and CsrA/FliW.

The analysis also suggested associations among poorly characterized flagellum-associated proteins, including YvyF with DUF327/DUF6470 proteins, MotE with P- and L-ring proteins, PilZ-domain proteins with FlaG, and FlaF with FlbT (Fig. 3d; Supplementary Fig. 2). These associations provide hypotheses for future functional studies rather than definitive evidence of direct interactions or shared function. In particular, associations represented by deeper branches of the hierarchical clustering may be less robust and should be interpreted in the context of their underlying pairwise similarity values.

### Ancestral reconstruction supports 50 ancestral flagellar components

Although 40 flagellar protein groups showed strong co-presence across bacterial genomes, co-presence alone does not establish ancestral presence. To support ancestry, a gene must also exhibit broad distribution across deeply diverged bacterial lineages in a pattern more consistent with vertical inheritance than with recent horizontal transfer. To investigate the evolutionary relationships among flagellar components, we developed a phylogenetic reconstruction approach that integrates information from 74 gene trees representing 37 broadly distributed flagellar protein groups (Fig. 4a). For each protein group, we constructed trees using full-length protein sequences and regions matched by profile HMMs. To reduce the influence of sequence divergence, horizontal gene transfer, and lineage-specific evolutionary events on individual gene trees, we integrated their phylogenetic signals at four taxonomic levels: family, order, class, and phylum.

**Figure 4:**
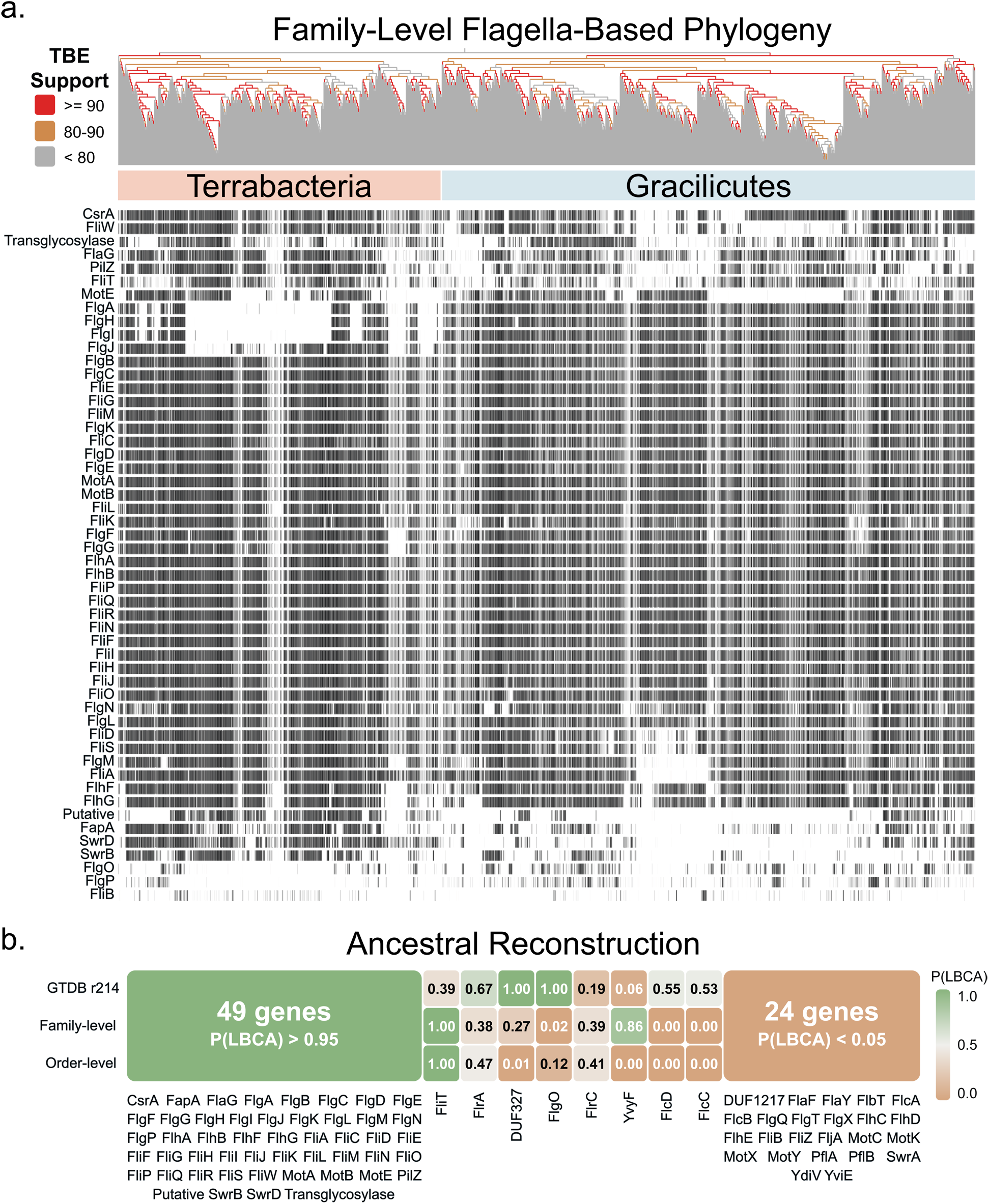
Phyletic distribution and ancestral reconstruction of flagellar proteins support the ancestral flagellar gene set. **a,** Presence/absence patterns for the 52 candidate ancestral flagellar protein groups are shown across the family-level flagella-based phylogeny displayed at the top, with branches colored by Transfer Bootstrap Expectation (TBE) support (red, ≥90%; orange, 80–90%; gray, <80%). The phylogeny separates two major clades, corresponding primarily to Terrabacteria (light red) and Gracilicutes (light blue). Each vertical column represents a bacterial taxonomic family, and black marks indicate the presence of a given gene within genomes belonging to that family. Families were normalized to equal horizontal width regardless of the number of sampled genomes, allowing patterns of gene presence and absence to be compared independently of uneven taxonomic sampling. The tool used to create this plot is available at https://www.flagelladb.org/phyletic-distribution-visualization. The underlying data can be further investigated at https://www.flagelladb.org/phyletic-distribution-table. **b,** Ancestral state reconstruction with PastML, performed using three phylogenetic frameworks: the GTDB r214 genome-level tree and two hybrid trees constructed using the family-level and order-level flagella-based phylogenies as backbones. Genes with unanimous high-confidence ancestral support (P(LBCA) > 0.95 in all three reconstructions) are collapsed into a single block (49 protein groups, left); genes with unanimous high-confidence support for absence (P(LBCA) < 0.05 in all three reconstructions) are collapsed into a second block (24 protein groups, right); both blocks list their member genes below. The remaining eight genes, for which the three reconstructions disagree, are shown individually, colored by their posterior probability of ancestral presence, P(LBCA).

Our phylogenetic reconstruction approach is based on the premise that taxonomic groups repeatedly clustering together across multiple flagellar gene trees are likely to share an evolutionary history. For each gene tree, we labeled sequences according to their taxonomic assignments and quantified the extent to which pairs of taxa clustered together across internal nodes, accounting for their representation within each clade. The resulting taxon-by-taxon similarity matrices were averaged across the 74 gene trees and converted into a distance matrix, from which we inferred a neighbor-joining tree. This procedure was performed independently at the family, order, class, and phylum levels. By integrating phylogenetic information across multiple flagellar components, this approach was intended to reduce dependence on individual gene histories and provide a summary of the evolutionary relationships reflected in the flagellar protein dataset.

To evaluate the robustness of our flagella-based phylogenies, we performed a bootstrap analysis by resampling the input protein groups and calculated branch support using Transfer Bootstrap Expectation (TBE). Furthermore, to benchmark the accuracy of the inferred topology, we compared our custom phylogenies against the GTDB r214 reference tree at each rank using four complementary metrics: Robinson-Foulds distance, Clustering Information Distance, quartet concordance, and Transfer Distance (Table S3). The family-level tree showed the strongest agreement with GTDB across all four metrics, with chance-corrected agreement values of 30.0%, 61.1%, 78.7%, and 58.6%, respectively. It also exhibited the highest branch support among the four taxonomic ranks with a median TBE of 92% and 55% of branches receiving TBE support of at least 90%. These results indicate that the family-level reconstruction captures substantial phylogenetic signal shared with the genome-scale reference, while retaining differences that may reflect the distinct evolutionary histories of flagellar proteins. We therefore used the family-level tree for visualizing the phyletic distribution of flagellar components (Fig. 4a).

The family-level flagella-based phylogeny resolved two major superclades corresponding predominantly to Gracilicutes and Terrabacteria (Fig. 4a), providing a natural framework for inferring ancestral gene presence and for placing the root of the tree between these two clades (TBE = 90%). The only main exception was a clade containing several Gracilicutes-affiliated genomes, from Spirochaetes, Fusobacteria, and Thermotogota, and several candidate phyla Lindowbacteria, Muirbacteria, Riflebacteria, Wallbacteria, UBP17, and T1Sed10-126, which grouped within the Terrabacteria superclade. Both the clade itself (TBE = 95%) and its placement within Terrabacteria (TBE = 86%) received substantial bootstrap support. This unexpected grouping could reflect horizontal transfer of flagellar genes between ancestral bacterial lineages, phylogenetic artifacts such as long-branch attraction, or a combination of these factors. Although the available data do not allow us to distinguish among these explanations, the broader separation of Gracilicutes and Terrabacteria remains a key feature of the reconstructed topology. To evaluate the ancestral presence of individual flagellar protein groups, we performed ancestral state reconstruction using PastML on three alternative phylogenetic frameworks: the GTDB r214 genome-scale bacterial phylogeny and two hybrid phylogenies constructed using our family-level and order-level flagella-based trees as backbones. In the hybrid trees, relationships within individual taxonomic groups were obtained from GTDB, allowing us to retain genome-level resolution while incorporating the alternative evolutionary relationships inferred from flagellar proteins. For each protein group, we estimated the probability of presence in the last bacterial common ancestor (LBCA) independently under all three frameworks (Fig. 4b; Table S4). Based on their presence across both Gracilicutes and Terrabacteria, we initially identified 52 flagellar protein groups as candidates for ancestral presence (Fig. 4a). Ancestral state reconstruction with PastML provided high-confidence support for 49 of these groups, each with a probability of presence in LBCA exceeding 0.95 in all three phylogenetic frameworks (Fig. 4b, Table S4). Of the remaining three candidates, FliT received strong support from both flagella-based phylogenies (P(LBCA) = 1.00), although its support using the GTDB tree was substantially lower (P(LBCA) = 0.39). Given its broad distribution across both major superclades, we consider FliT an additional likely ancestral component. FlgO showed conflicting support across the three frameworks (P(LBCA) = 1.00, 0.02, and 0.12 for GTDB, family-level, and order-level trees, respectively), whereas FliB received little support for ancestral presence. Together, these results identify 50 likely ancestral flagellar protein groups, including 49 supported at high confidence across all three phylogenetic reconstructions, more than doubling the previously estimated ancestral set of 24 components.

In addition to reconstructing ancestral components, the phyletic mapping also identified several protein groups with restricted taxonomic distributions (Supplementary Fig. 3), consistent with their emergence or recruitment during subsequent flagellar evolution. These included YvyF, DUF327, and DUF6470 in the common ancestor of Terrabacteria, and FlrA and FlrC in the common ancestor of Gracilicutes. Some of these groups, such as DUF327, FlcC, and FlcD, received support in individual reconstructions (Fig. 4b), but their restricted distributions suggest that this reflects the placement of the GTDB r214 root close to Spirochaetota rather than ancestral presence. The remaining flagellum-associated protein groups showed restricted distributions, consistent with auxiliary components acquired later in flagellar evolution.

### Gene Retention and Absence for Ancestral Core Components

We next examined retention of the 50 inferred ancestral flagellar protein groups in extant bacterial genomes. Among genomes encoding at least 25 of these components, many retained between 41 and 44 groups (Fig. 5a), indicating substantial conservation of the ancestral repertoire. Notably, 12 Bacillota_C genomes retained the complete set of 50 ancestral protein groups. Genomes encoding at least 49 ancestral components were found in both Gracilicutes and Terrabacteria, indicating that highly complex flagellar systems are not restricted to a single bacterial lineage and supporting the presence of a similarly complex apparatus before the divergence of the two major superclades (Supplementary Fig. 4).

**Figure 5:**
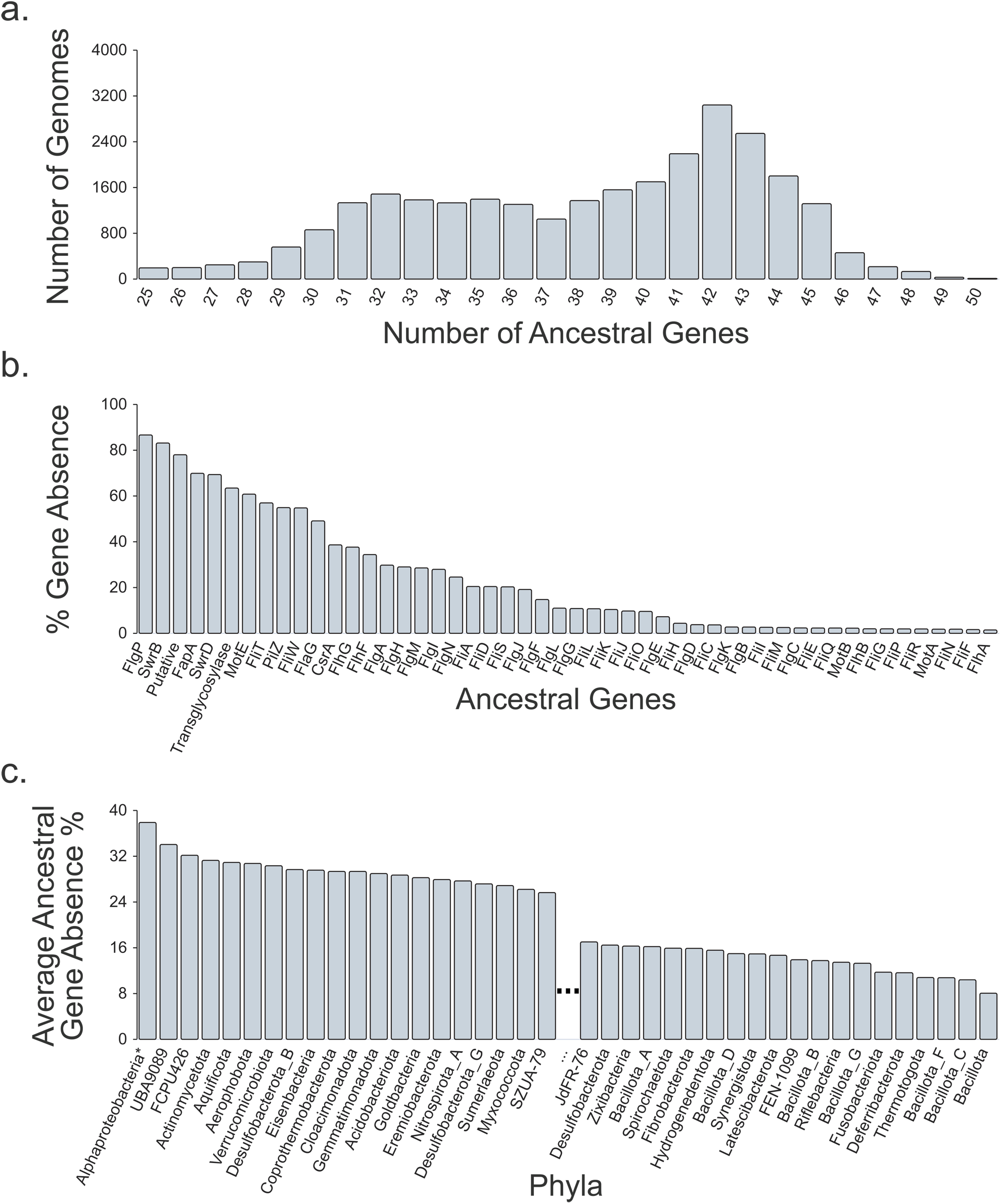
Retention and absence patterns of ancestral flagellar proteins. **a,** Distribution of retained ancestral flagellar proteins across genomes encoding at least 25 of the 50 inferred ancestral protein groups. **b,** Gene-specific absence frequencies for each ancestral flagellar protein across the same genome set, shown as the percentage of genomes in which a given gene is absent. **c,** Total ancestral gene absence percentages across bacterial phyla, calculated using the same genome set as in panels a and b. For visualization, only the top 20 and bottom 20 phyla (restricted to phyla with at least 5 members) are shown. To minimize biases associated with differences in cell-envelope architecture, outer membrane-associated proteins (*FlgA*, *FlgH*, *FlgI*, and *FlgJ*), often co-absent in monoderm genomes, were excluded from this analysis. Because of its large size and phylogenetic diversity, *Pseudomonadota* are shown at the class level.

We next quantified absence frequencies for individual ancestral protein groups across the same genome set (Fig. 5b). Proteins involved in core structural, export, and motility functions, including FlhA, FlhB, MotA, MotB, FliQ, FliR, FliP, FliC, FliG, and FliE, were absent from only ∼2–5% of genomes. The rare absence of FliC is consistent with its use as a simple predictor of flagellar motility ^30^, although other flagellar components may still be retained in its absence. In contrast, regulatory and accessory components showed substantially higher absence frequencies. The retention pattern is consistent with differences in functional constraints: genes required for assembly, export, and motility are strongly retained due to their indispensable roles. In contrast, genes with regulatory or accessory functions are more frequently absent, perhaps reflecting lineage-specific selective pressures. Notably, proteins containing the transglycosylase domain we included in this work showed poor co-presence with other ancestral flagellar components (Fig. 3d) and were absent from more than 60% of genomes (Fig. 5b), despite their broad phyletic distribution (Fig. 4a). This pattern suggests that these proteins are likely ancestral, but may have roles that are not specific to bacterial flagella. The same may apply to broadly acting regulators, such as CsrA and PilZ-domain proteins. Although the inferred ancestral presence of these components makes secondary loss a plausible explanation for their absence in extant lineages, absence from our dataset does not necessarily indicate evolutionary loss, as it may also reflect incomplete genome assemblies, sequence divergence, or limitations of protein detection.

To assess broader phylogenetic patterns, we next quantified total gene absence percentages at the phylum level (Fig. 5c). Because *Pseudomonadota* is both highly represented and phylogenetically diverse, its classes were analyzed separately. To avoid bias associated with cell-envelope architecture, outer membrane-associated proteins (FlgA, FlgH, FlgI, and FlgJ), which tend to be missing together from monoderm genomes, were excluded from this comparison, and lineages represented by fewer than five genomes were omitted. The lowest average absence frequencies (∼8-10%) were observed in several Bacillota-associated lineages (*Bacillota*, *Bacillota_C*, and *Bacillota_F*), as well as *Fusobacteriota*, *Thermotogota*, *Riflebacterota*, and *Deferribacterota*. In contrast, *Alphaproteobacteria* showed the highest average absence frequency (37.91%), followed by the candidate phyla *UBA9089* and *FCPU426*, and *Actinomycetota*, each with average absence frequencies exceeding 30%. The lineage-specific differences indicate substantial variation among bacterial lineages in the retention of ancestral flagellar components.

Together, these results support a model in which the LBCA possessed a complex flagellar system, whereas the repertoires of extant bacteria have been shaped by differential retention and secondary loss of ancestral components. Proteins involved in core structural, export, and motility functions are generally strongly retained, while regulatory and accessory components exhibit more variable distributions. Thus, much of the variation in flagellar composition among extant bacterial lineages can be explained by differential retention of an already complex ancestral repertoire.

## DISCUSSION

The bacterial flagellum has long served as a model for the evolution of complex molecular systems ^13,26,46,47^. Although previous studies have provided evidence that the last bacterial common ancestor (LBCA) possessed a flagellum ^28^, its ancestral composition has remained uncertain. Earlier reconstructions identified 24 ancestral flagellar components ^26^, substantially fewer than those found in extant flagella. Here, by combining large-scale homolog detection, co-presence analysis, phyletic distribution, and probabilistic ancestral state reconstruction, we identify 49 flagellar protein groups with high-confidence support for their presence in the LBCA, together with one additional likely ancestral component. Our reconstructed repertoire of 50 protein groups (Fig. 6) more than doubles previous estimates and indicates that the LBCA possessed a flagellar apparatus substantially more complex than previously recognized. We acknowledge that ancestral reconstructions frequently infer ancestors more complex than many of their descendants, as reported for the last universal and last eukaryotic common ancestors ^48,49^. This tendency could partly reflect methodological artifacts biased by horizontal gene transfer, uneven taxonomic sampling, or inaccurate assumptions about rates of gene gain and loss. Alternatively, the last common ancestor of a domain need not have been primitive, since the LBCA was itself the product of earlier evolution. Although we combined a qualitative assessment of gene presence across both bacterial superclades with ancestral state reconstructions on three different phylogenies, our data do not allow us to fully distinguish between these two possibilities.

**Figure 6.**
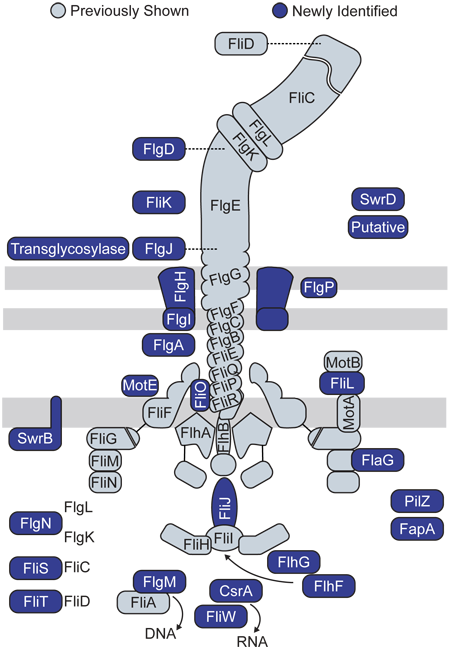
Reconstruction of the ancestral bacterial flagellum. Schematic overview of the flagellar apparatus inferred to have been present in the LBCA. The reconstruction comprises 49 protein groups supported as ancestral with P(LBCA) > 0.95 across three phylogenetic frameworks, together with FliT as an additional likely ancestral component. Proteins previously identified as ancestral components are shown in light blue, whereas additional ancestral components identified in this study are shown in dark blue. The reconstructed repertoire includes the principal structural, export, motor, and assembly components required for a functional flagellum. An extended representation incorporating major lineage-specific flagellum-associated proteins is provided in Supplementary Fig. 5.

Our findings indicate that a complex and functionally complete flagellar apparatus, containing all major modules required for its assembly and rotation, was already present in the LBCA and that subsequent flagellar evolution involved differential retention, secondary loss, and lineage-specific innovation. However, the evolutionary events that led to the emergence of this complex molecular machine before the LBCA remain only partly understood. Previous studies have identified homologous relationships among flagellar proteins and between flagellar components and proteins functioning in other cellular systems, including ATP synthases and protein secretion systems ^13,50^. These relationships, together with evidence for duplication and diversification of flagellar structural proteins ^26^, provide important clues to the evolutionary origins of individual components and suggest that recruitment and modification of preexisting proteins contributed to flagellar evolution. Nevertheless, they do not establish the sequence of events through which these components became integrated into a functional, rotating flagellar apparatus. Thus, while our reconstruction substantially expands the inferred ancestral repertoire, elucidating the evolutionary processes that produced this complex system before the LBCA remains an important challenge.

The presence of a functionally complete flagellum in the LBCA may have provided early bacterial lineages with an important ecological advantage, enhancing their ability to explore diverse environments and contributing to the long-term persistence of flagellated descendants. Subsequent flagellar evolution appears to have been dominated primarily by secondary loss and lineage-specific innovation. This is consistent with independent evidence that flagellar motility has been lost far more often than gained across bacteria ^30^ and that flagella in Actinobacteria underwent extensive degenerative loss from a common flagellated ancestor ^44^. During this phase, many extant flagellated bacteria retained over 40 flagellar components (Fig. 5a), indicating that much of the ancestral complexity was preserved. However, gene retention patterns show that this preservation was not uniform. Structural, export-associated, and motility-associated components are retained at much higher frequencies (Fig. 5b), consistent with strong selective pressure to preserve the essential architecture and function of the flagellum. In contrast, regulatory and accessory components are absent more frequently, possibly because their functions are more readily compensated or rewired in different cellular and ecological contexts. For example, FlaY performs a FliD-like filament-capping function in *Caulobacter crescentus* despite a distinct fold ^51^, and flagellar gene expression is controlled by FlhDC in enteric bacteria but by unrelated master regulators, such as FlrA in *Vibrio*, in other lineages ^52,53^. Lastly, differences in gene absence patterns across lineages (Fig. 5c) are consistent with lineage-specific trajectories of flagellar reduction and remodeling.

Our findings suggest that the functional organization of the flagellum was largely established before the diversification of major bacterial lineages. Co-presence analysis identified strongly associated protein groups corresponding to known functional modules, including the T3SS-related module, stator, rod, hook, filament, and several regulatory circuits (Fig. 3, Supplementary Fig. 2). Each core module remains recognizable across deeply diverged bacterial lineages, consistent with the early establishment and subsequent conservation of much of the flagellum’s functional organization. This suggests that the ancestral flagellum was not only complex, but already organized into semi-independent functional units that could be differentially modified ^45^.

Our findings also highlight a broader methodological point. Deep evolutionary reconstructions are often limited by the difficulty of detecting highly diverged homologs and distinguishing true gene absence from limitations of sequence-based searches. To maximize sensitivity, we performed sequence searches using relaxed E-value thresholds and primarily relied on phylogenetic clustering to distinguish target flagellar protein groups from other homologous sequences. Nevertheless, we identified cases in which previously characterized orthologs showed unexpectedly low scores against the HMM profiles or iterative sequence profiles used in our initial searches. For FliF, FliH, FlrC and MotB, we addressed this by rerunning the pipeline using alternative orthologs from lineages where the initial search failed to recover clear candidates. These examples demonstrate that even sensitive homology detection pipelines may fail to detect highly diverged homologs, potentially leading to overestimation of gene absence. Similarly, very short or unusually long protein sequences may occasionally cluster within retrieved homologous clades during high-throughput analysis. In addition, our flagella-based phylogenies are phenetic summaries of the flagellar gene trees rather than species trees. This allows them to reflect the shared history of flagellar genes, including horizontal transfer, without assuming that these genes followed the species tree. However, because they are built at the family and order levels, they do not resolve relationships among individual genomes within these groups. For ancestral state reconstruction, we combined them with the GTDB reference tree, which provided the relationships within each group, and also used the GTDB tree on its own. Therefore, although the retention and absence patterns reported here provide a broad evolutionary framework, they remain the product of a large-scale automated pipeline and could be further refined through additional targeted curation. To facilitate further refinement through community feedback, we have made our dataset publicly available through flagelladb.org, allowing researchers to explore and evaluate flagellar protein distribution across bacterial lineages.

Together, our findings provide a substantially revised reconstruction of the bacterial flagellum in the last bacterial common ancestor. Rather than a relatively simple precursor, the LBCA likely possessed a complex, functionally complete flagellar apparatus comprising at least 49 high-confidence ancestral protein groups and one additional likely ancestral component. The diversity of flagellar systems observed in extant bacteria is consistent with extensive differential retention, secondary loss, and lineage-specific modification of this ancestral repertoire. Although the evolutionary processes that gave rise to the flagellum before the LBCA remain incompletely understood, our reconstruction establishes a more comprehensive framework for investigating its early evolution and subsequent diversification.

## METHODS

### Compilation and curation of the reference flagellar protein set

We compiled a comprehensive reference set of flagellum-associated proteins using the KEGG pathway map02040 (“Flagellar assembly”), which contains 54 proteins, as the initial source. To identify additional components not represented in KEGG, we assembled a reference set of 251 bacterial genomes selected to provide broad taxonomic coverage, include experimentally characterized flagellar systems, and prioritize genome quality and representation in the MiST database ^36^. We searched these genomes using profile hidden Markov models (HMMs) corresponding to the initial protein set and examined the genomic neighborhoods of identified flagellar genes to detect additional proteins consistently associated with flagellar loci. Candidate proteins were further identified through a literature survey emphasizing experimentally characterized components involved in flagellar assembly, structure, function, and regulation. Chemotaxis proteins were excluded because they constitute a distinct signal-transduction system rather than components of the flagellar apparatus.

These approaches yielded an initial collection of 95 flagellum-associated proteins. We subsequently used sequence similarity, domain architecture, and available functional annotations to identify homologous proteins representing the same functional group and consolidated these, together with synonymous gene names, into a nonredundant set of 81 orthologous groups (Tables 1 and 2). Here, orthologous groups are defined operationally as groups of homologous proteins associated with the same flagellar function and may include lineage-specific paralogs. This curated set served as the reference for subsequent large-scale homology searches across bacterial genomes.

### Homology searches and sequence retrieval

For each flagellar gene, we used HMMs from the KOfamKOALA ^38^ and Pfam ^39^ databases when available, and from TIGRFAMs ^40^ and NCBIfam ^41^ for YvyF and FliB, respectively. HMM searches were performed with HMMER v3.3.2 ^54^ using the “-E 1000” option. When no HMMs were available, we used previously annotated protein sequences as queries in iterative profile searches performed with MMseqs2 ^55^ using the “-e 1000 --e-profile 10 -s 7.5 --num-iterations 5” options. MMseqs2 searches were used for FlaY, FlcA, FlcB, FlcC, FlcD, FlgX, FljA, FlrA, MotC, MotK, MotX, PflA, PflB, SwrA, SwrB, and YdiV, as well as for the reruns of FliF, FliH, FlrC, and MotB. All other protein groups were searched with HMMER. All database searches were conducted against GTDB release 214 ^37^ proteomes, covering 80,789 representative bacterial genomes. For most protein groups, we retrieved the top 50,000 sequence matches ranked by search score. This cutoff was chosen empirically because our initial analyses suggested that about one third of genomes in GTDB r214 contain flagellar genes, and we therefore aimed to exceed the expected size of the orthologous group to maximize sensitivity. For multicopy proteins such as FliC, MotA, and MotB, we retrieved the top 100,000 hits. Because these proteins frequently occur in multiple copies per genome, the top 50,000 hits consisted almost entirely of orthologous group members, leaving too few background sequences to delineate the group boundary and missing some members.

### Filtering of Inferred Orthologous Groups

#### Sequence Alignment and Gene Tree Inference

For each set of retrieved sequences, we generated a multiple sequence alignment (MSA) using FAMSA v2.2.2 ^56^ with default settings. Alignment columns containing more than 90% gaps were trimmed using trimAl v1.4 ^57^ with the “-gt 0.1” option, and gene trees were inferred from the trimmed alignments using FastTree v2.1.11 ^58^ with default settings (JTT substitution model with CAT approximation of rate heterogeneity). For searches based on HMM models, we performed this workflow both on the full retrieved sequences and on the regions matched by the HMM model. For each flagellar component, we then manually selected the tree that provided the clearest separation of the orthologous group from background noise.

#### Ordering the Gene Tree and MSA

To facilitate visual identification of orthologous groups, tree leaves were ordered by HMMER profile-HMM scores or MMseqs2 bit scores, depending on the search method (hereafter search scores), using a recursive algorithm. First, each tree was rooted on the single lowest-scoring sequence. The non-target sequences surrounding each orthologous group often included related paralogous families or background matches. We then traversed the tree from the root through all internal nodes and, for each pair of sister clades, calculated the average search score. When necessary, sister clades were rearranged so that the clade with the higher average score appeared on the left. This ensured that high-scoring sequences were concentrated on the left side of the tree. The resulting tree order was also applied to the corresponding MSAs. We then plotted search scores, rescaled to range from 0 to 1, together with sequence rank in the search results to identify tree regions enriched in high-scoring sequences that also ranked highly in the original search output. Because ordering only rearranges sister clades without changing the tree topology, it served as a visual aid rather than an independent filtering criterion and did not determine which sequences were assigned to an orthologous group.

#### Identifying Gene-Gene Neighbors

We then compared the sequences in each gene search output against up to 100,000 sequences from the outputs of the other flagellar gene searches. For each sequence, we identified genes located on the same genome, contig, and strand, and classified a gene as a neighbor if it was located within 500 bp of the query sequence. We then aggregated this neighborhood information into windows of 50 consecutive sequences along the ordered tree and plotted it as a line plot to highlight tree regions associated with specific neighboring genes.

#### Identification Criteria

To identify sequences belonging to each flagellar orthologous group, we evaluated four complementary criteria (Supplementary Fig. 1): (i) clustering within phylogenetic trees, (ii) search scores reflecting sequence similarity to the query, (iii) genomic association with other flagellar genes, and (iv) conserved sequence patterns identified through visual inspection of multiple sequence alignments using the Jalview Overview Window. Sequences forming phylogenetic clades supported by these combined criteria were assigned to the corresponding orthologous groups and retained for downstream analyses. Importantly, individual sequences were not required to satisfy every criterion independently. For example, sequences with relatively low similarity scores or without detectable flagellar gene neighbors could still be included when they clustered within a clade supported by the remaining evidence. This approach was designed to maximize recovery of divergent flagellar proteins while reducing the inclusion of unrelated sequences or homologs belonging to different functional groups. The underlying sequence alignments, phylogenetic trees, search results, and associated visualizations are available through our Zenodo repository, with an interactive view of each orthologous group at https://www.flagelladb.org/transparency/orthologous-groups.

#### Benchmarking against the manually curated reference set

To estimate the sensitivity of the bacteria-wide detection pipeline, we compared it against the manually curated reference set (251 representative genomes) at the 238 genome assemblies present in both. For each of 80 matched flagellar protein groups, we pooled every individual protein/gene ID reported by each dataset and computed a true-positive rate (TPR) in both directions, treating the curated set as reference (TP = IDs found in both datasets, FN = curated IDs the pipeline missed) and treating the pipeline as reference (FN = pipeline IDs the curated set did not confirm), with TPR = TP / (TP + FN) in each case. A genome/gene comparison was excluded from this calculation whenever any of the pipeline’s hits for that gene in that genome had a GTDB id that failed to translate to an NCBI accession, since this reflects an ID-mapping gap rather than a genuine detection difference. Every remaining ID-level mismatch between the two datasets was checked against NCBI’s most recent protein record, and mismatches confirmed to be stale or withdrawn accessions were removed from the false-negative count in their respective direction rather than counted as either a hit or a miss. TPR was left undefined for genes with zero comparable reference IDs, which occurred either because the curated set had no entries for that gene at all (FljA) or because the genome carrying its only curated entry was excluded by the translation-failure rule above (YdiV). Per-gene TP, FN, and TPR values for both directions, along with median TPR across all matched genes, are provided in Table S2.

### Gene Co-presence Analysis

We analyzed 55 widely distributed flagellar protein groups across the 80,789 representative bacterial genomes in GTDB r214. For a given gene pair, we calculated the Jaccard similarity index as number of genomes in which both genes were detected divided by number of genomes in which at least one of the two genes was detected (Fig. 3a).

To reduce the influence of uneven taxonomic sampling, we applied family-level inverse weighting, which corrects for heavily sampled lineages while retaining resolution among closely related genomes. Each genome was inversely weighted according to the number of genomes sampled from its bacterial family, such that genomes from highly represented families contributed less individually than genomes from sparsely represented families. For each gene pair, the weighted contributions of individual genomes containing both genes were summed and divided by the summed weighted contributions of genomes containing at least one of the two genes (Fig. 3b). This produced a family-balanced weighted Jaccard similarity index (J_w_) for each gene pair.

Pairwise weighted Jaccard similarity scores were converted to distances by subtracting weighted Jaccard indexes from one (Fig. 3c). Genes were then grouped using complete-linkage hierarchical clustering, and the clustered similarity matrix was visualized as a heatmap (Fig. 3d).

### Constructing Flagella-based Phylogeny

To reconstruct phylogenetic relationships from flagellar proteins, we developed an approach that integrates phylogenetic information across individual gene trees rather than concatenating protein sequences into a single alignment. Our custom algorithm quantifies how frequently pairs of taxonomic groups cluster together within gene trees and combines these associations to infer a phylogeny representing their shared signal. We applied this procedure independently at four taxonomic ranks (family, order, class, and phylum), producing a separate flagella-based phylogeny at each rank.

#### Selection of flagellar gene trees

For this analysis, we extracted the inferred orthologous groups for each flagellar gene from the manually selected clades described above. For each gene, we used the full-length-sequence tree and the HMM-matched-region tree as two separate inputs, increasing the amount of data while keeping each gene’s contribution to the analysis equal. We selected 37 broadly distributed flagellar protein groups for phylogenetic analysis. These groups included FlgB, FlgC, FlgD, FlgE, FlgK, FlgL, FlhA, FlhB, FliC, FliD, FliE, FliF, FliG, FliH, FliI, FliK, FliM, FliN, FliP, FliQ, FliR, FliS, MotA, MotB, FliO, FliJ, FliL, FliW, FlaG, FlgM, FlgN, FlhG, FlhF, FlgJ, FlgG, FlgF, and Transglycosylase. For each group, we extracted sequences belonging to the previously identified orthologous group and used two phylogenetic trees: one inferred from full-length protein sequences and another from HMM-matched regions. The resulting dataset comprised 74 gene trees representing 37 flagellar protein groups. Both trees associated with each protein group were included in the subsequent analysis, maintaining equal representation of the selected groups.

#### Taxonomic labeling and node scoring

For each taxonomic rank, every leaf of each gene tree was labeled with its GTDB r214 taxonomic assignment at that rank. For example, a leaf sequence from an Escherichia coli genome would be labeled Enterobacteriaceae when building the family-rank tree, but Pseudomonadota when building the phylum-rank tree, since the same leaf takes a broader or narrower label depending on which rank is being analyzed. Because many genomes belong to the same family or phylum, this labeling routinely produces multiple leaves in the same tree that share an identical taxon label, for example several sequences from different Enterobacteriaceae genomes all labeled Enterobacteriaceae within a single family-rank tree. To reduce the influence of underrepresented taxonomic lineages, we removed taxa represented by fewer than two sequences.

For a given rank, all 74 flagellar protein group trees were labeled and traversed from the root toward the terminal leaves, recording which taxa were present under each internal node and how many sequences each contributed.

#### Calculation of the similarity matrix per gene tree

To calculate the taxon-by-taxon similarity matrix for a given gene tree, we quantified, for each internal node, the co-clustering of every pair of taxa present there. To do this, we first scaled each taxon’s count at the node by dividing it by that taxon’s total sequence count across the whole tree, so the scaled value reflects the proportion of that taxon’s sequences captured under this node rather than its raw abundance. This scaling reflects how concentrated, rather than how numerous, a taxon’s sequences are at a given node: a node capturing most of a taxon’s sequences represents a stronger association than one capturing only a small fraction of them, regardless of the raw sequence counts involved. It also prevents large, heavily sampled taxa, whose sequences can be scattered across many parts of the tree simply due to their size or occasional horizontal transfer, from generating false evolutionary signal due to their high abundance. For example, if a taxon is represented by 40 sequences across the whole tree and 30 of them fall under a given node, its scaled value at that node is 30/40, or 0.75. For every pair of taxa present at a node, we then multiplied their two scaled proportions to obtain that node’s co-clustering score for the pair, so a score near 1 indicates that nearly all of both taxa’s sequences cluster together under this node, while a low score indicates that only a small fraction of one or both taxa’s sequences are present there. We summed these node-level scores across every internal node of the tree where the pair co-occurred, producing one similarity value per taxon pair, and repeating this across all taxon pairs produced the tree’s full similarity matrix. We then normalized this matrix by its own highest value, so that every gene tree’s matrix was rescaled onto the same 0-1 scale before being combined with the other trees.

#### Combining similarity matrices across gene trees

To calculate the overall similarity between taxon pairs across all of the 74 gene trees, we averaged their normalized similarity scores across trees. However, due to uneven coverage of individual genes across lineages, we divided each pair’s summed similarity by the number of trees in which both taxa were actually present, rather than by a fixed count of 74, so the average reflected only the trees where the pair could be meaningfully compared.

Taxa with zero similarity to all other taxa, which never clustered with another taxon in any gene tree, were removed because their placement could not be inferred. Separately, we also tracked how many of the 74 trees each individual taxon appeared in on its own and removed any taxon present in fewer than 80% of trees from the final matrix entirely. This threshold was chosen to retain as much taxonomic coverage as possible while still excluding taxa represented in too few trees for their similarity values to be reliable.

#### Construction of the distance matrix and neighbor-joining tree

Finally, the combined similarity matrix was converted into a distance matrix by subtracting each similarity value from one. In this distance matrix, taxon pairs with high co-representation across flagellar gene trees had shorter distances, whereas taxon pairs with low co-representation had longer distances. The resulting distance matrix was then used to construct a neighbor-joining tree. The trees generated at family, order, class, and phylum levels each represent a flagella-based phylogeny that summarizes the collective phylogenetic signal shared across the analyzed flagellar genes at varying levels of resolution. Because the combined matrix summarizes how often taxa co-occur across flagellar gene trees rather than modeling vertical descent, these trees are phenetic summaries of the flagellar gene trees and not species trees.

#### Branch support

For each rank, we assessed how robust the inferred phylogenies were to gene sampling using a gene-resampling bootstrap, while keeping the set of taxa across bootstrap replicates identical to the inferred tree. We resampled the 37 flagellar genes with replacement, randomly drawing 37 genes each time and including both trees (full-length and HMM-matched-region) associated with every sampled gene, then repeated the full matrix-construction and neighbor-joining pipeline described above on each resampled gene set. This was repeated 100 times per rank, producing 100 replicate distance matrices and trees. We then calculated Transfer Bootstrap Expectation (TBE) support for each branch by comparing it against the 100 replicate trees.

### Benchmarking Tree Topology Against GTDB Taxonomy

#### Preparing trees for comparison

We compared each rank-level flagella-based tree to the GTDB r214 bac120 reference tree. To ensure proper comparison, the GTDB reference tree was collapsed to the family, order, class, and phylum level, and both the GTDB and custom-phylogeny trees were then pruned to the taxon set they shared. Because the flagella-based neighbor-joining trees are unrooted, all metrics were computed based on unrooted topologies.

#### Topology comparison metrics

We used four metrics to quantify topological agreement, computed in Python (ete3 package) for Robinson-Foulds distance and quartet concordance, and in R (TreeDist) for Clustering Information Distance and Transfer Distance. Normalized Robinson-Foulds distance measures the fraction of bipartitions that differ between the two trees. Clustering Information Distance measures split disagreement while weighting by how much phylogenetic information each split carries, giving partial credit for near-miss splits. Quartet concordance measures the fraction of resolved four-taxon subtrees on which the two trees agree; we estimated it by Monte Carlo sampling of 2,000,000 quartets rather than exact enumeration. Transfer Distance measures the minimum number of taxa that would need to move for a branch in one tree to match its counterpart in the other tree.

#### Assessing statistical significance

We assessed each metric’s significance using a permutation test, which measures how well the metric would score by chance alone if the flagella tree’s shape had no real relationship to GTDB’s topology. This lets the actual agreement between the two real trees be judged against that random baseline, rather than assumed to be meaningful on its own. To do this, the flagella tree’s tip labels were randomly shuffled 499 times while the GTDB tree was kept fixed, effectively generating 499 randomized versions of the flagella tree, and each metric was recalculated on every such pairing to build an empirical null distribution and p-value. Robinson-Foulds distance, Clustering Information Distance, and Transfer Distance were each chance-corrected by rescaling against their own empirical null mean from these permutations, while quartet concordance was instead rescaled based on its theoretical minimum: since any four taxa can be resolved in exactly three possible ways, two unrelated trees would be expected to agree on one-third of quartets by chance alone, a floor that closely matched the empirical permutation nulls where checked.

Across all four metrics and branch-support measures, the family-level tree showed the strongest agreement with the GTDB topology and the highest TBE support (Table S3), and was therefore used for the qualitative visualization of ancestral flagellar gene presence and absence (Fig. 4a). The family-level and order-level trees, the two best-performing ranks, together with the GTDB r214 reference tree, were used as three alternative phylogenetic frameworks for the quantitative ancestral state reconstruction described below.

### Ancestral State Reconstruction

#### Generating the input trees

For the ancestral state reconstruction, we used the GTDB r214 bacterial genome-level tree and two hybrid phylogenies constructed from our family- and order-level custom phylogenies. The GTDB reference tree only required minor reformatting of its internal node labels for standard tree-parsing compatibility. Its leaf names and topology remained unchanged. Each custom flagella-based phylogeny, however, was used to create a hybrid tree that followed the backbone of the flagella-based hierarchy, while the relationships within each group instead followed the reference GTDB tree. This approach allowed ancestral state reconstruction to be performed using genome-level gene presence/absence data under alternative phylogenetic backbones, enabling us to assess the sensitivity of ancestral gene assignments to differences in deeper tree topology.

#### Ancestral state reconstruction with PastML

For each of the three trees, we encoded every flagellar protein group as a binary presence/absence character (1 if the genome had at least one homologous sequence for that gene, 0 otherwise) and provided this table to PastML as input, matched and reordered to that tree’s own genome order. Ancestral states were reconstructed independently for each of the three trees using PastML’s marginal posterior probabilities approximation (MPPA) method under the F81 substitution model, inferring a posterior probability of flagellar protein group presence, P(LBCA), at the root node of each tree (Fig. 4b).

#### Inferring Ancestral Gene Set

For each flagellar protein group, we classified its ancestral state as unanimous, high-confidence presence at the LBCA if P(LBCA) exceeded 0.95 in all three reconstructions, and as unanimous, high-confidence absence if P(LBCA) fell below 0.05 in all three. Protein groups for which the three reconstructions did not agree at this threshold were classified as disputed and reported individually rather than assigned a single call (Fig. 4b). These quantitative posterior probabilities were evaluated alongside an independent qualitative assessment of each group’s phyletic distribution, specifically, whether it was broadly present across both major bacterial superclades, to arrive at the final set of inferred ancestral protein groups. The full posterior probabilities for all genes across all three reconstructions, including calls at additional confidence thresholds (0.5, 0.7, and 0.9) and the final classification of each group, are provided in Table S4.

### Calculating Gene Retention and Absence Percentages

#### Calculating ancestral gene retention per genome

To evaluate the retention of ancestral flagellar components in extant bacteria, we restricted the analysis to genomes encoding at least 25 of these 50 inferred ancestral protein groups. For each genome in this dataset, the total number of present ancestral protein groups was plotted as a bar plot in Figure 5a. Here, retention is defined as the detection of at least one identified member, irrespective of copy number.

#### Calculating gene-specific absence percentages

To quantify the frequency of absence of individual ancestral flagellar components, we analyzed the same dataset of genomes containing at least 25 of the 50 inferred ancestral protein groups. For each protein group, the absence percentage was calculated as the number of genomes in which no member of that group was detected, divided by the total number of genomes analyzed, multiplied by 100. Protein groups were then ranked by their absence percentages to identify components that were most frequently retained or absent across extant bacterial genomes (Fig. 5b).

#### Calculating lineage-specific gene absence

To measure the frequency of ancestral flagellar protein group absence across broader taxonomic groups, we calculated average absence percentages at the phylum level, which summarizes broad patterns across major bacterial lineages rather than inferring ancestral states. To prevent biases originating from differences in cell-envelope architecture, outer membrane-associated protein groups (FlgA, FlgH, FlgI, and FlgJ), whose joint absence is common in monoderm genomes, were excluded, leaving a subset of 46 ancestral protein groups for analysis. Because of its large size and substantial phylogenetic diversity, the phylum *Pseudomonadota* was analyzed at the class level. Lineages represented by fewer than five genomes were excluded to minimize sampling artifacts. For each retained taxonomic group, we calculated the absence percentage of each of the 46 ancestral protein groups across its constituent genomes. These percentages were then averaged to obtain an overall absence percentage for that taxonomic group. Lineages were ranked by this average absence metric, and the 20 lineages with the highest and lowest values were visualized for comparison (Fig. 5c).

## Data Availability

All MSAs, phylogenetic trees, inferred flagellar orthologous groups, and intermediate files and visualizations generated by our computational pipeline are available through our Zenodo repository. Custom tools necessary to generate Figures 3–5 are available in our database at flagelladb.org.

Zenodo: https://doi.org/10.5281/zenodo.22266385

## Code Availability

The custom code used in this study is available through our GitHub repository. GitHub: https://github.com/Jouline-Lab/LBCA_Flagellum

## Acknowledgements

This work was supported by the National Institutes of Health grants R35GM131760 to I.B.Z. and R35GM131783 to D.B.K. and by Medical Research Council grant MR/Z504385/1 to M.B. M.E. acknowledges funding from the European Research Council (ERC) under the European Union’s Horizon 2020 research and innovation program (grant agreement No. 864971) and from the Max Planck Society as a Max Planck Fellow.

## Author contribution

Conceptualization: I.B.Z. and M.E.; bioinformatic analyses: B.S. and E.P.A.; investigation and interpretation: B.S., E.P.A., M.B., D.B.K., M.E., and I.B.Z.; writing: B.S., E.P.A., and I.B.Z.; supervision: I.B.Z.

