## Supplemental Figures for "The last bacterial common ancestor encoded the components of a functionally complete flagellum"

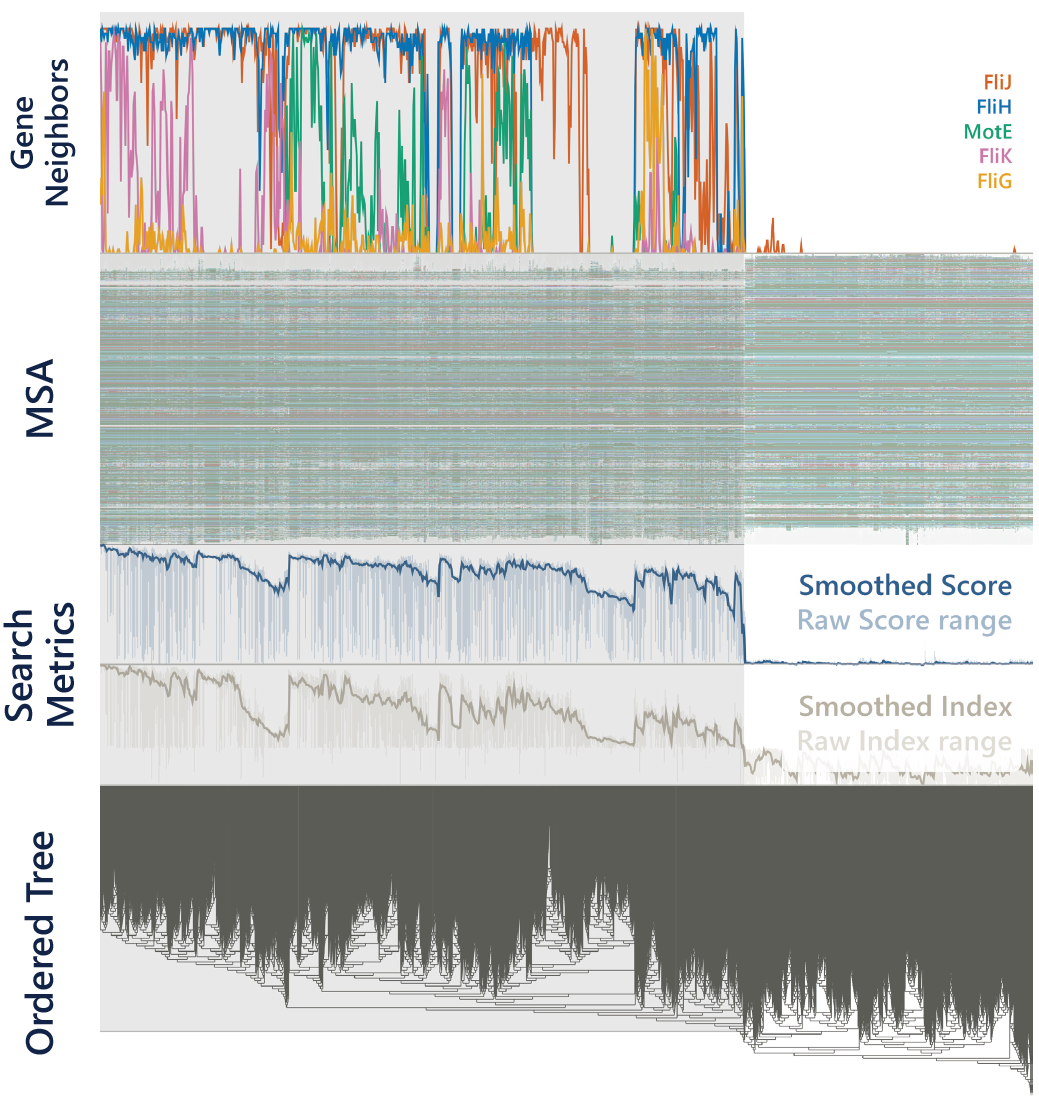

**Figure S1. Example application of the orthologous-group identification pipeline to the flagellar ATPase FliI.** The four tracks correspond to the four criteria used to define orthologous groups: the normalized search score and search-rank index, neighboring flagellar genes, the tree-ordered MSA, and the search-score-ordered gene tree itself. The inferred orthologous group, highlighted in grey, was defined as the clade whose sequences consistently show higher search scores, frequent flagellar-gene neighbors (FliJ, FliH, MotE, FliK, and FliG), and differential conservation patterns in the MSA. Because FliI was searched with a profile HMM, search scores here are HMMER profile-HMM scores, rescaled to range from 0 to 1. Neighbor lines show the fraction of each window of 50 consecutive sequences with that gene encoded within 500 bp. Search-score and rank-index lines are smoothed, with shaded bands spanning the unsmoothed values. An interactive version of this figure, for FliI and for each of the other orthologous groups, is available at https://www.flagelladb.org/transparency/orthologous-groups?group=FliI.

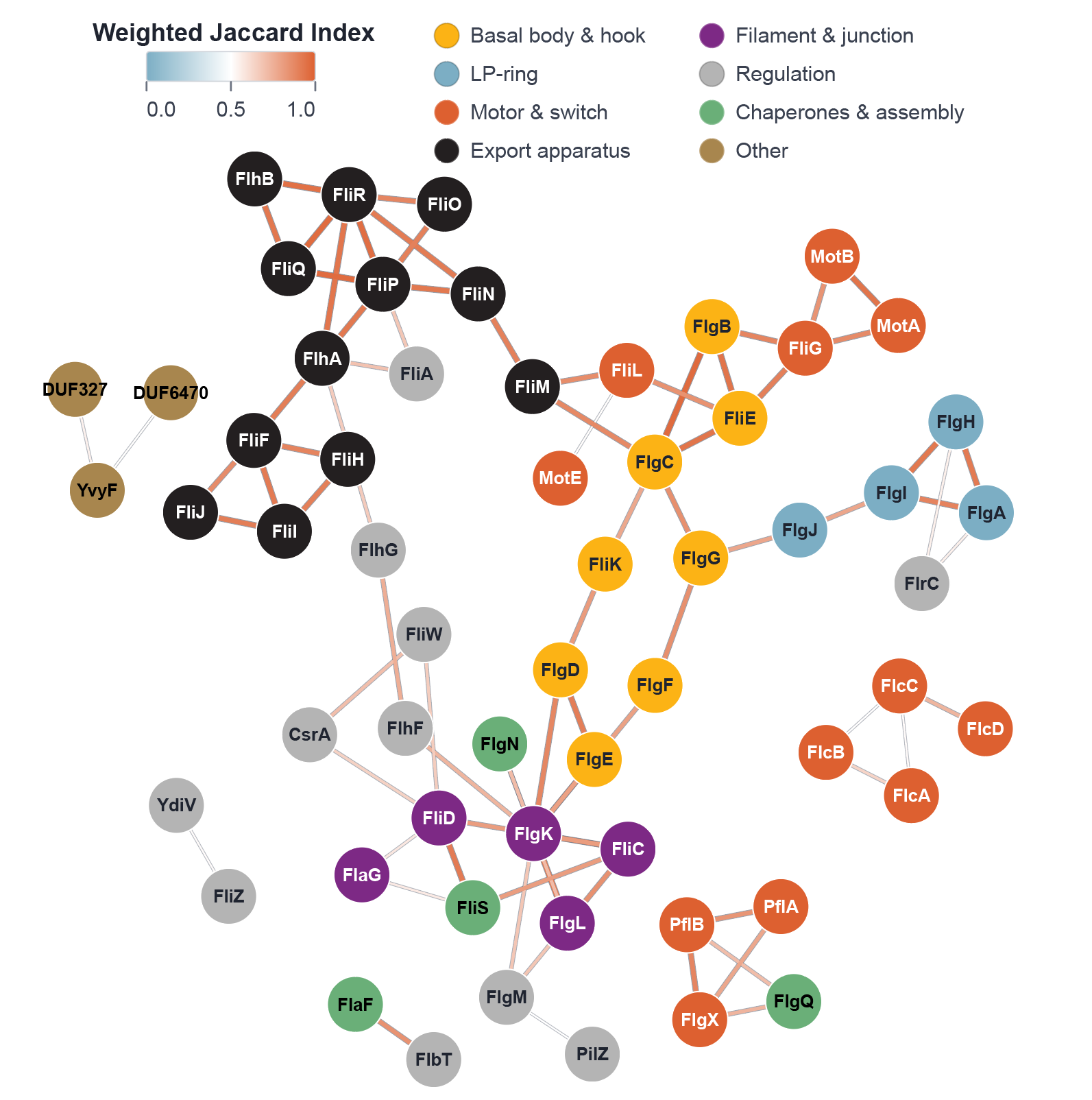

**Figure S2: Network representation of family-weighted co-occurrence among flagellar genes.** Force-directed network visualization of the pairwise family-weighted Jaccard similarities (Figure 3d). Each node represents a flagellar gene and is colored according to its functional category. Edges connect gene pairs with a family-weighted Jaccard similarity >0.5, and edge color indicates similarity strength according to the scale shown. To highlight the strongest co-occurrence relationships while reducing network complexity, only the two highest-scoring connections were retained for each gene. Because this selection is made independently for each gene, some genes have more than two edges, and genes with a single retained edge may appear visually prominent without being strongly connected. Node positions are determined by the layout algorithm and do not represent quantitative similarity. Genes lacking any retained connections were omitted from the network, leaving 59 of the 81 flagellar orthologous groups analyzed. The complete, unfiltered similarity data are available at https://www.flagelladb.org/gene-correlation.

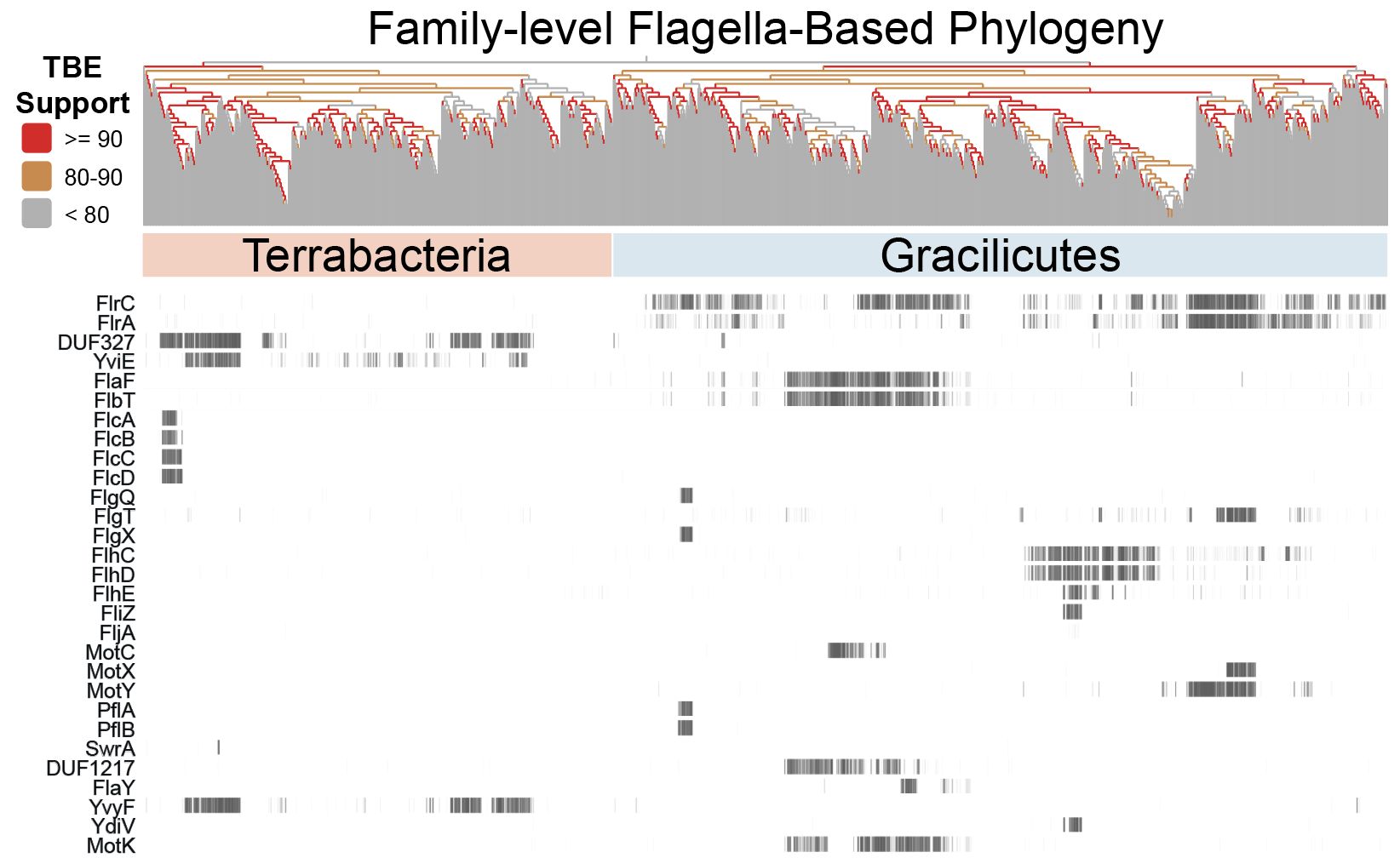

**Figure S3: Phyletic distribution of lineage-specific flagellar proteins**

Presence/absence patterns for the 29 flagellar protein groups with lineage-specific distribution are shown across the same family-level flagella-based phylogeny as in Figure 4a, with branches colored by Transfer Bootstrap Expectation (TBE) support (red, ≥90%; orange, 80-90%; gray, <80%). The phylogeny separates two major clades, corresponding primarily to Terrabacteria (left) and Gracilicutes (right). Each vertical column represents a bacterial taxonomic family, and black marks indicate the presence of a given gene within genomes belonging to that family. In contrast to the broadly distributed ancestral gene set shown in Figure 4a, these genes show restricted, lineage-specific distributions, consistent with acquisition after the divergence of the major bacterial superclades.

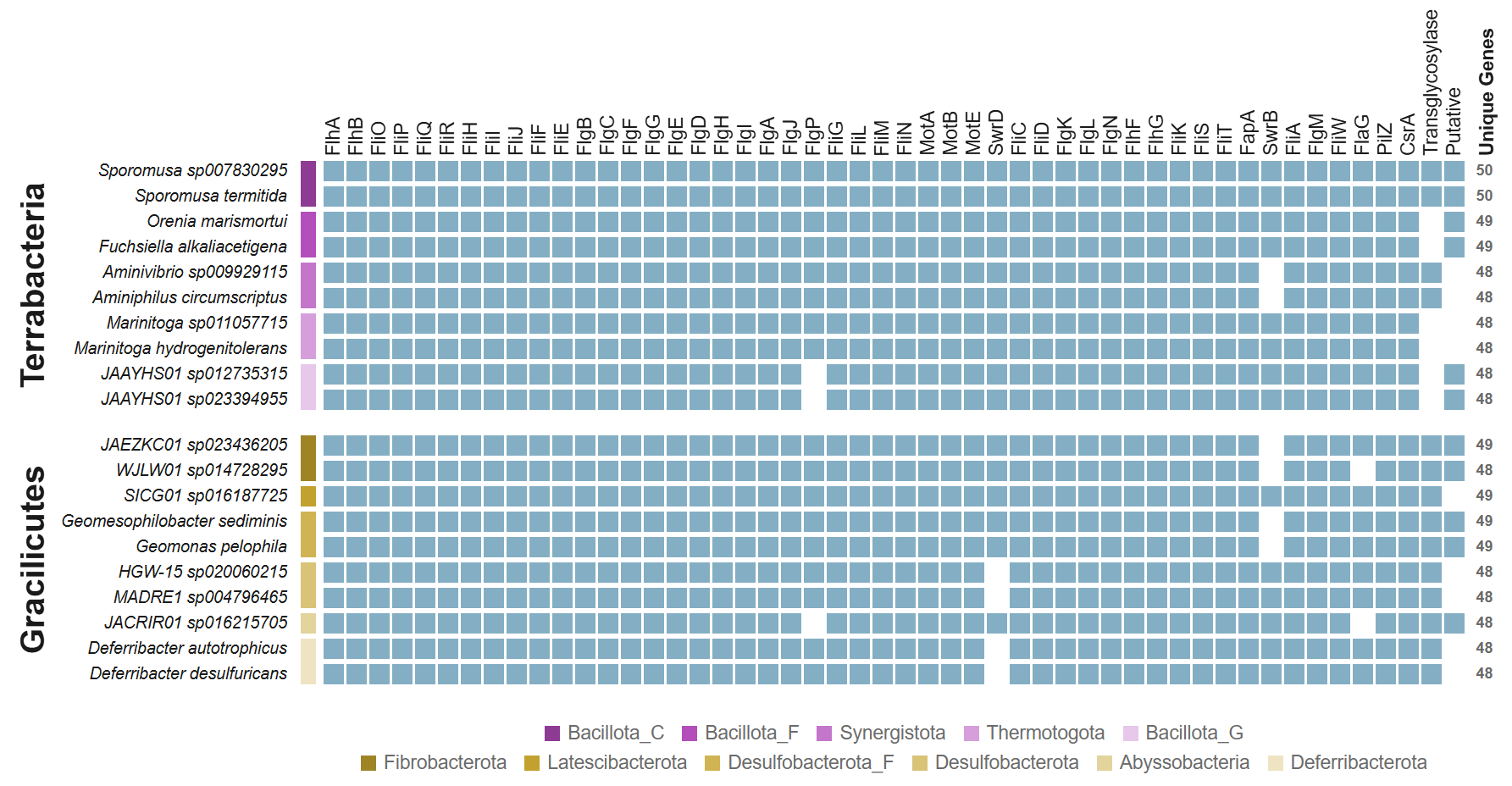

**Figure S4: Species with the highest number of retained ancestral flagellar components in both Terrabacteria and Gracilicutes.**

Presence and absence of the 50 inferred ancestral flagellar genes, shown as columns, for the ten genomes with the most complete ancestral gene sets in each superclade, shown as rows and colored by taxonomic phylum as indicated in the legend. Filled cells indicate gene presence and blank cells indicate absence. The rightmost column reports the total number of ancestral genes detected in each genome. Genomes retaining 48-50 of the 50 ancestral genes are found in both superclades, indicating that highly complex flagellar systems are not restricted to a single bacterial lineage.

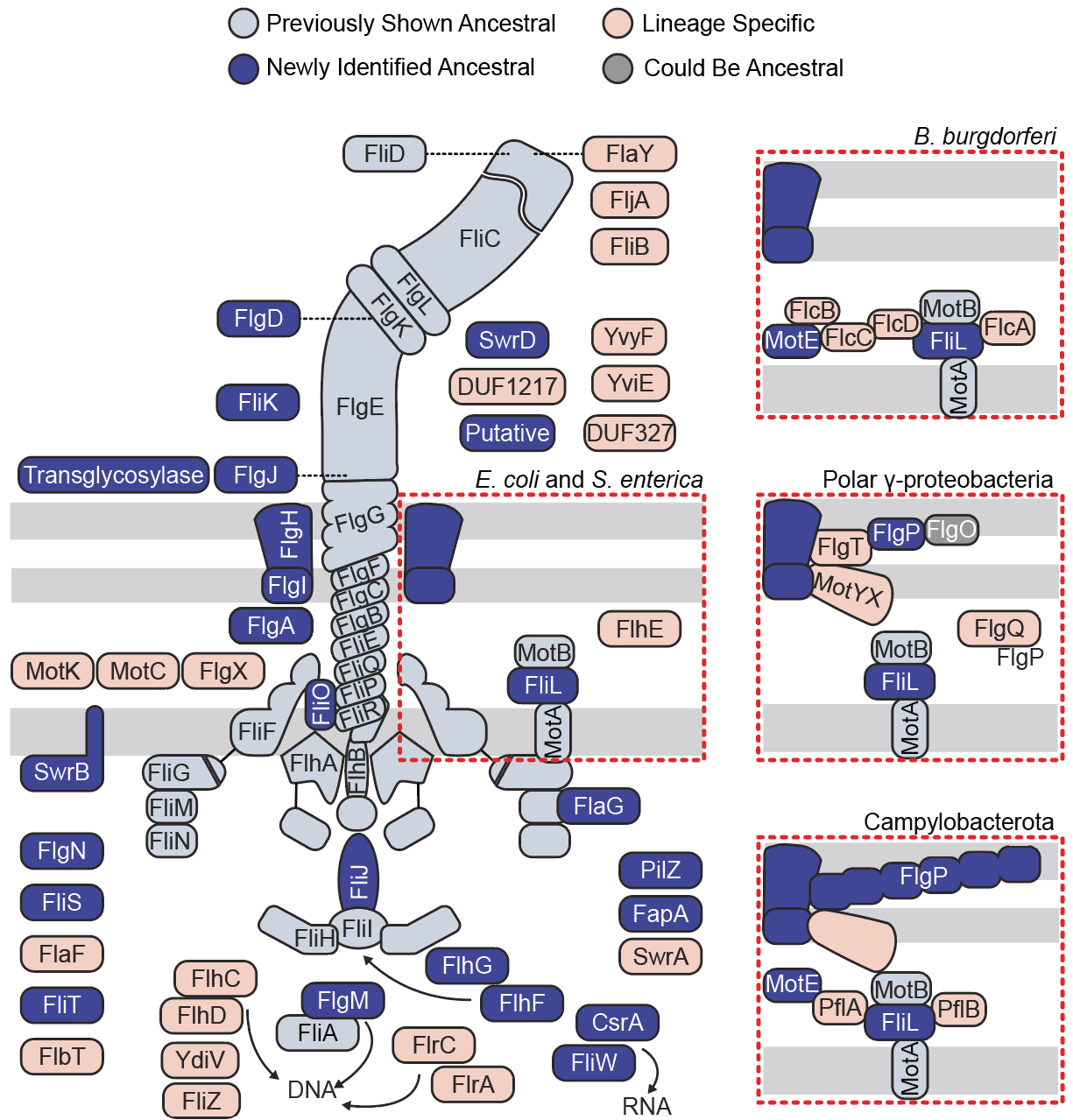

**Figure S5. Reconstructed ancestral bacterial flagellum and lineage-specific flagellum-associated proteins.** Schematic representation of the flagellar system inferred to have been present in the Last Bacterial Common Ancestor (LBCA). Components previously proposed to be ancestral are shown in light blue, whereas additional ancestral proteins identified in this study are shown in dark blue. Proteins inferred to have been acquired later during the diversification of bacterial lineages are shown in pink. FlgO is shown in grey to indicate its uncertain ancestral status, as it is distributed across both superclades but its ancestral presence is supported only by the GTDB r214 reconstruction. Insets illustrate representative lineage-specific additions associated with the flagella of *Escherichia coli* and *Salmonella enterica*, Spirochaetota (*Borrelia burgdorferi*), polar γ-proteobacteria, and Campylobacterota.
